# Understanding site choice and competition through the adaptive dynamics of microbial settling strategy in a chemostat

**DOI:** 10.1101/2020.07.13.189662

**Authors:** Yuhua Cai, Stefan A. H. Geritz

## Abstract

To understand the choice and competition of sites in nature, we consider an ecological environment in a chemostat consisting of a polymorphic microbial population that can float in the fluid or settle down on the wall of the chemostat. For the transition of a microbe from its floating state to its settled state at a particular settling rate involving the choice and competition of sites on the wall, we consider three different mechanisms: (*i*) *unimolecular-Bourgeois settling*, i.e., floaters land freely on the wall, but in an occupied site, the owner keeps the site (Bourgeois behaviour); (*ii*) *unimolecular-anti-Bourgeois settling*, i.e., floaters land freely on the wall, but in an occupied site, the intruder gets the site (anti-Bourgeois behaviour); (*iii*) *bimolecular settling*, i.e., floaters land only on the vacant sites of the wall. Employing the framework of adaptive dynamics, we study the evolution of the settling rate with different settling mechanisms and investigate how physical operating conditions affect the evolutionary dynamics. Our results indicate that settling mechanisms and physical operating conditions have significant influences on the direction of evolution and the diversity of phenotypes. (1) For constant nutrient input, theoretical analysis shows that the population is always monomorphic during the long-term evolution. Notably, the direction of evolution depends on the settling mechanisms and physical operating conditions, which further determines the composition of the population. Moreover, we find two exciting transformations of types of Pairwise Invasibility Plots, which are the *gradual transformation* and the *bang-bang transformation*. (2) For periodic nutrient input, numerical analysis reveals that evolutionary coexistence is possible, and the population eventually maintains dimorphism. Remarkably, for all three settling mechanisms, the long-term evolution leads to one of the two coexisting species settle down totally on the wall if the input is low-frequency but float entirely in the fluid if the input becomes high-frequency.

## 1. Introduction

Site (or habitat) choice is a vital decision in the lives of many organisms. For instance, oviposition site choice is an essential component of the life history of mosquitoes. Laboratory and field studies have shown that oviposition site choice by most mosquito species is not random (see Yoshioka et al. 2012 and references therein). A female mosquito seeking oviposition sites must often balance the risk of predation and competition to their progeny (see, e.g., Blaustein 1999; Kershenbaum et al. 2012). For birds, the risk of nest predation influences habitat settlement decisions. Predator removal experiments show that birds respond to the presence of predators by altering settlement decisions (Fontaine and Martin 2006). The additional example concerning the choice of sites in ecology is the refuge (or shelter) that allows a species to be protected against unfavourable conditions (Hawkins et al. 1993; Cowlishaw 1997; Sih 1997; Berryman et al. 2006). For instance, in coral reef community, a prey obtains protection from predation by hiding in a reef hole (see, e.g., Hixon and Beets 1993); in a chemostat, microbes escape the implication of dilution operation by settling down on the wall of the chemostat (ref to the attachment phenomenon in microbial growth which has stimulated considerable research, see, e.g., Baltzis and Fredrickson 1983; Freter et al. 1983; Smith and Waltman 1995; Pilyugin and Waltman 1999; Stemmons and Smith 2000; Ballyk et al. 2008; Harmand et al. 2017).

When the amount of vacant sites is abundant, individuals can quickly settle. However, habitable sites are restricted in many situations. For instance, there are only limited sites on the wall of a chemostat for microbes to adhesion (see, e.g., Ballyk et al. 2008). When sites are limited, individuals have to compete with the owner in an occupied site. Generically, there are two population dynamical outcomes of such a competition event: either the owner keeps the site (Bourgeois behaviour) or the intruder gets the site (anti-Bourgeois behaviour) (ref to the contest dynamics of the Hawk-Dove model in Evolutionary Game Theory, see, e.g., Maynard Smith 1982; Mesterton-Gibbons and Sherratt 2014; Takeuchi 2018).

The choice and competition of sites is a complicated process that may involve the physical conditions (e.g., temperature and humidity) and the ecological properties (e.g., predator, prey and food). The mechanism is not yet well understood at the individual level, even for the examples mentioned above. Such behaviour makes the population model more nonlinear and hence affect the ecological-evolutionary dynamics.

Since the mechanism of choice and competition of sites is rather complicated and may be altered for different organisms, we cannot develop a single mathematical model that captures every aspect of the mechanism. Therefore, in this paper, the mechanistic modelling with simplifications is employed to gain an insight into the mechanism.

To understand site choice and competition in nature, we consider a stage-structured microbial population in a chemostat model. The microbial population consists of floating individuals (in the fluid) and settled individuals (on the wall). For the transition of a microbial individual from its floating state to its settled state at a particular settling rate involving the choice and competition of sites, we consider three different mechanisms: (*i*) *unimolecular-Bourgeois settling*, i.e., floaters land freely on the wall, but in an occupied site, the owner keeps the site; (*ii*) *unimolecular-anti-Bourgeois settling*, i.e., floaters land freely on the wall, but in an occupied site, the intruder gets the site; (*iii*) *bimolecular settling*, i.e., floaters land only on the vacant sites of the wall.

The purpose of this paper is to provide a mathematical model for understanding the mechanism of choice and competition of sites and investigate its impacts on the ecological-evolutionary dynamics. To do so, we study the evolution of microbial settling rate with different settling mechanisms in a chemostat utilising the framework of adaptive dynamics (see, e.g., Metz et al. 1992, 1996; Geritz et al. 1997, 1998, 1999).

Chemostat is a device for culturing microorganisms that provides excellent utility in tracking their ecological-evolutionary dynamics (Gresham and Hong 2015). In a simple chemostat model, two physical operating conditions are under control: the concentration of nutrient input and the dilution rate. The effects of the two physical operating conditions on ecological dynamics have attracted much attention (see, e.g., Hale and Somolinos 1983; Butler et al. 1985; Yang and Freedman 1991; Ruan 1993; Smith 1997; Ruan and He 1998; Caraballo et al. 2015, 2018; Nguyen et al. 2020). In this paper, we investigate how the physical operating conditions affect the evolution of microbial population with different settling mechanisms.

The organisation of the paper is as follows. In Sect. 2, we derive an ecological model in a chemostat consisting of a polymorphic microbial population who can float in the fluid or settle down on the wall of the chemostat. In particular, three different settling mechanisms are proposed, which concern the choice and competition of sites on the wall. In Sect. 3, we establish some analysis results of the ecological dynamics with a single microbe type, which concern the survival analysis in the virgin environment, the permanence of the system, the existence and uniqueness of a positive equilibrium, and the global stability of the positive equilibrium. In Sect. 4, the evolution of the settling rate is analysed by using the framework of adaptive dynamics. In particular, we classify different evolutionary outcomes for the three settling mechanisms in Sect. 4.2. Our results show that the population is always monomorphic during the long-term evolution, whatever the settling mechanisms and physical operating conditions change. In Sect. 5, we apply numerical analysis to the evolution with non-equilibrium resident dynamics, which reveals the possibility of evolutionary coexistence and the way of coexistence in a fluctuating environment. Finally, we conclude our results in Sect. 6. All proofs are given in the Appendices.

The definitions of state variables and model parameters of the ecological model are given in Tab. 1, in which only nonnegative values are meaningful. Unless specifically requested, we omit the argument *t* in all time-related variables. Throughout this paper, the dot denotes differentiation with respect to time variable *t*, the superscript ′ denotes differentiation with respect to its associated argument of a function, and the superscript T means transpose of a vector or a matrix.

**Tab. 1.**
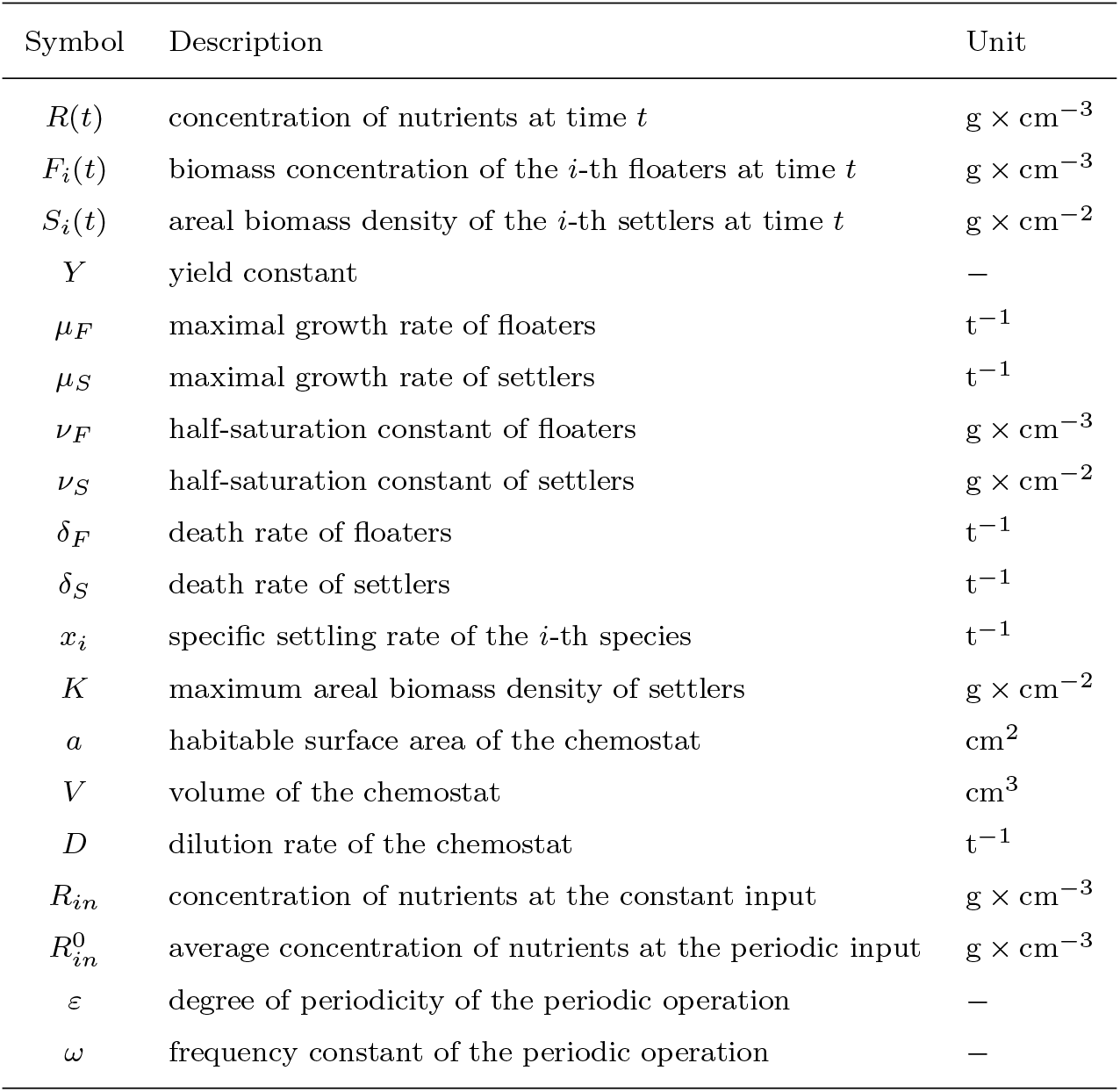
Definitions of state variables and model parameters.

## 2. Derivation of the ecological model

Consider an ecological environment in a chemostat consisting of a polymorphic microbial population that can float in the fluid or settles down on the wall of the chemostat. Different species is charactered by different settling rate. Thus, we divide the *i*-th species with settling rate *x_i_* into floaters *F_i_* and settlers *S_i_*. Both floaters and settlers consume a common nutrient *R* and then reproduce their floating offsprings at a constant ratio *Y*. Generally, *Y* is called the yield (Microbiology) or the efficiency (Ecology). Since the organism is the same in our model, one anticipates that the yield constants are equal whether the microbe is in its floating state or its settled state (see, e.g., Pilyugin and Waltman 1999; Stemmons and Smith 2000; Ballyk et al. 2008). Freter et al. (1983) assumed that the specific growth rate of a microbe in a biofilm model is the same whether the microbe is in its planktonic state (i.e., the floating state) or its wall-attached state (i.e., the settled state) for a given value of the nutrient concentration. Subsequent evidence from work on biofilms suggests that the different growth rates of the microbe in its different states are more realistic (see Stemmons and Smith 2000 and references therein). In this paper, we consider a large class of growth rates *f_F_*(*R*) and *f_S_*(*R*) for floaters and settlers respectively, which requires continuity in *R* with *f_F_* (0) = *f_S_*(0) = 0, and monotonicity in *R*, i.e., 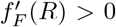 and 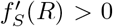 for all *R* > 0. A common choice is the Monod growth functional response:

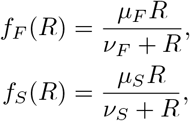

where *μ_F_* (resp. *μ_S_*) is called the maximal growth rate and *ν_F_* (resp. *ν_S_*) is the half-saturation (or Michaelis-Menten) constant for floaters (resp. settlers) (Monod 1949; Smith and Waltman 1995).

Due to the limitation of habitable sites on the wall of the chemostat, the model assumes a maximum areal biomass density *K* for microbes that can settle down on the wall.

Let ***x*** = (*x*_1_,…, *x_k_*), ***F*** = (*F*_1_,…, *F_k_*), and ***S*** = (*S*_1_,…, *S_k_*). A floater of the *i*-th species lands on the wall at rate *g_i_*(***x, S***) that depends on the corresponding settling rate and the ratio of candidate sites (thus it is involved in the biomass density of settlers). Because of competition in occupied sites, only a part of them successfully transits into the settled group at rate *h_i_*(***x, F, S***). The formulations of *g_i_* and *h_i_* will be given with interpretations under different mechanisms later.

Once a floater settles down on the wall of the chemostat, it can not slough off into the fluid again. Floaters will be removed by washout at rate *D*, compete-caused death in occupied sites at rate *g_i_*(***x, S***) – *h_i_*(***x, F, S***), and natural death at rate *δ_F_*, while settlers only undergo the natural death at rate *δ_S_*. Fig. 1 shows the transition of the microbial individual in its different states.

**Fig. 1.**
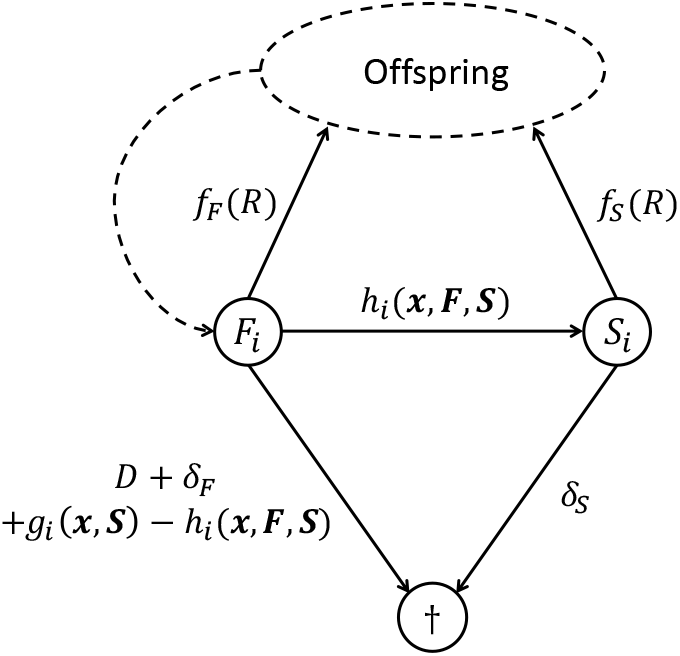
Diagram of floating-settled transition, in which the absorbing state † means removal due to washout, or compete-caused death in occupied sites, or natural death.

Assume that the culture vessel is well mixed with volume *V* and habitable surface area *a*. Moreover, all operating conditions are strictly controlled. Nutrients enter the chemostat at rate *DR_in_* and unused nutrients leave at rate *DR*(*t*), where *D* is the dilution rate and *R_in_* is the concentration of nutrient at the input.

With the above assumptions, we obtain the following system of differential equations for the polymorphic population dynamics in the chemostat,

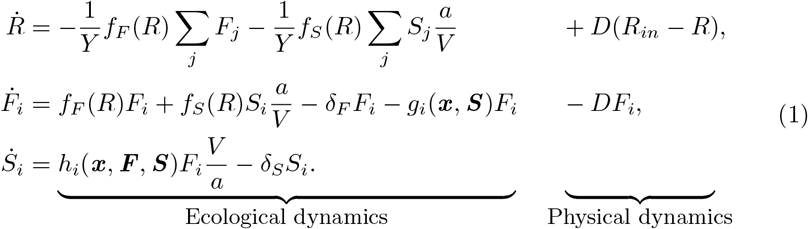

Without loss of generality, (1) can be simplified by setting 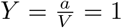. Therefore, we will apply this simplification to all subsequent equations.

To explain how a floater transits into the settled group, we consider the following three mechanisms:

i. *unimolecular-Bourgeois settling* (denote as *uB*-settling): a floater lands freely whatever the site is vacant or occupied. In vacant sites, landing implies settling. In each occupied site, the owner defeats the intruder and successfully guards the territory, i.e., the owner keeps the site (Bourgeois behaviour). The individual-level behaviour of such a mechanism would be

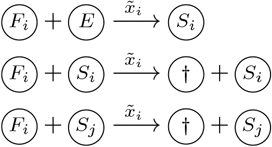

where 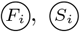, and 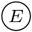 denote a floating individual of type *i*, a settled individual of type *i*, and a vacant site, respectively, and 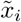 is the particular rate of *uB*-settling for the floaters of type *i*.
ii. *unimolecular-anti-Bourgeois settling* (denote as *uaB*-settling): it is similar to the *uB*-settling that a floater lands freely whatever the site is vacant or occupied. In each occupied site, however, the intruder kills the owner and then takeovers the territory, i.e., the intruder gets the site (anti-Bourgeois behaviour). The individual-level behaviour becomes

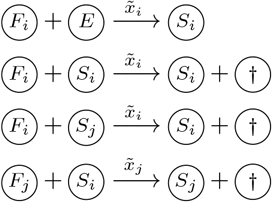

where 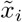 is the particular rate of *uaB*-settling for the floaters of type *i*.
iii. *bimolecular settling* (denote as *b*-settling): different from the previous two mechanisms, floaters only choose vacant sites to land. The corresponding individual-level behaviour is

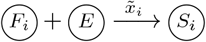

where 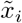 is the particular rate of *b*-settling for the floaters of type *i*.

For the continuous-time change in biomass, we thus have the term

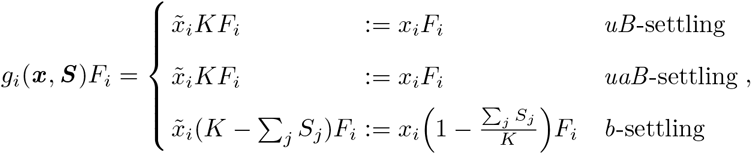

and the term

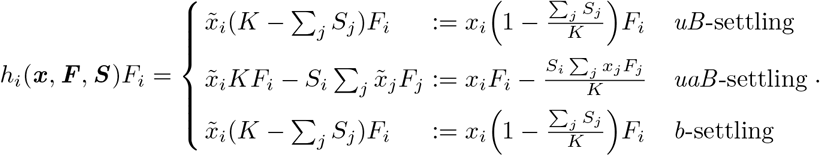

Here we have applied the change of variables, i.e., 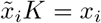. Notice the difference of the unit of 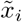 compared to that of *x_i_*.

If settlers do not reproduce floating offsprings, the wall serves only as a spatial refuge for the population. Further, the model we proposed here is a specific case of Pilyugin and Waltman (1999, (6.1)). In addition, if all microbial individuals are only in their floating state, our model reduces to the classical chemostat model (Novick and Szilard 1950).

Next is to investigate the evolution of the settling rate under different settling mechanisms and its ecological and evolutionary responses to changes in the physical operating conditions, i.e., the concentration of nutrient input *R_in_* and the dilution rate *D*.

## 3. Ecological dynamics with a single microbe type

When only a single microbe type is present, the ecological dynamics described by (1) are given by

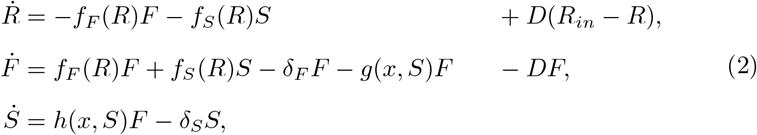

where

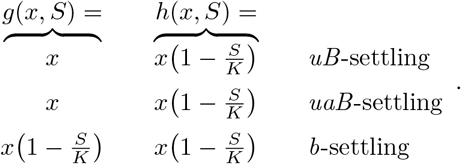

Notice that, in the monomorphic population dynamics, the function *h* is independent of *F*. As we see, the term *g*, as well as the term *h*, are identical for both the case of *uB*-settling and the case of *uaB*-settling. Hence, the ecological dynamics are the same whatever the monomorphic population adopts the *uB*-settling or the uaB-settling. Next, the analysis of the ecological dynamics of (2) will be divided into the case of *uB-/uaB*-settling and the case of *b*-settling, respectively.

### 3.1. Survival analysis in the virgin environment

When the system (2) is at the washout equilibrium *E*_0_ = (*R_in_*, 0,0), an initially rare microbial population with settling rate *x* is viable if and only if

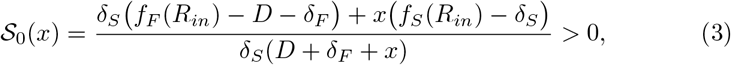

whatever the settling mechanism is. 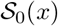 is obtained by calculating – 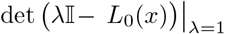 where 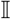 is a unit matrix and *L*_0_(*x*) is the next-generation matrix of the population dynamics of x in the virgin environment (Metz and Leimar 2011, and see Appendix A for details).

Now we introduce three quantities that play a central role in the analysis of ecological-evolutionary dynamics of (2). Let

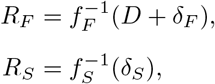

if they exist, otherwise we denote them as +∞. In fact, *R_F_* is the *break-even concentration* of the microbial population when floaters present alone, and *R_s_* is *break-even concentration* of the microbial population when settlers present alone (Smith and Waltman 1995). Notice that the break-even concentration *R_F_* is a function of *D*. Let

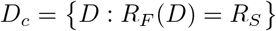

as the critical dilution rate. Moreover, the dilution operation with values

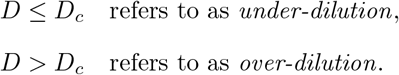

Here *D_c_* isn’t necessarily positive. If *D_c_* < 0, then we only consider the situation of *D* > 0. It is not difficult to check that *D* ≤ *D_c_* is equivalent to *R_F_* ≤ *R_S_* and that *D* > *D_c_* is equivalent to *R_F_* > *R_S_*.

The sign of 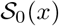 is dominated by the term *f_F_*(*R_in_*) – *D* – *δ_F_* that is the intrinsic growth rate of the microbial population when floaters present alone in the virgin environment, and the term *f_S_*(*R_in_*) – *δ_S_* that is the intrinsic growth rate of the microbial population when settlers present alone in the virgin environment. By the two intrinsic growth rates together with the quantity *D_c_*, there are four possible combinations to guarantee that the condition (3) holds:

I *f_F_*(*R_in_*) > *D* + *δ_F_* and *f_S_*(*R_in_*) ≤ *δ_S_*,
II *f_F_*(*R_in_*) ≤ *D* + *δ_F_* and *f_S_*(*R_in_*) > *δ_S_*,
III.a *f_F_*(*R_in_*) > *D* + *δ_F_* and *f_S_*(*R_in_*) > *δ_S_* with *D* ≤ *D_c_*,
III.b *f_F_*(*R_in_*) > *D* + *δ_F_* and *f_S_*(*R_in_*) > *δ_S_* with *D* > *D_c_*.

If the physical operating conditions (*D, R_in_*) satisfy case I, the chemostat is conducive to the survival of floaters. Moreover, the population of *x* is viable if its settling rate is smaller than *x_e_* where 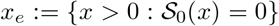. Conversely, if (*D, R_in_*) satisfy case II, settlers become superior to floaters. And the population of *x* will be viable if its settling rate is larger than *x_e_*. For cases III.a and III.b, 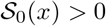 for all *x* ≥ 0 which means that the population with any settling rate x is always viable in the virgin environment. Particularly, settlers have a stronger nutrient uptake ability than that of floaters if the chemostat performs an under-dilution operation, while the situation is reversed if the chemostat performs an over-dilution operation. Fig. 2a shows the viable region in terms of *D, R_in_* and *x*. The population of *x* is easy to survive if the chemostat has a low-valued dilution rate and a high-valued concentration of nutrient input. Fig. 2b gives a cross-section through Fig. 2a at *x* = 2 to show the above classification, in which the boundary *R_F_*(*D*) approaches infinity if *D* tends to *D_v_* and the invasion boundary 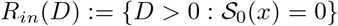 approaches infinity if *D* tends to *D_e_*. Here we would like to highlight the fact that no population can survive without nutrient input, i.e., *R_in_* = 0 and/or *D* = 0.

**Fig. 2.**
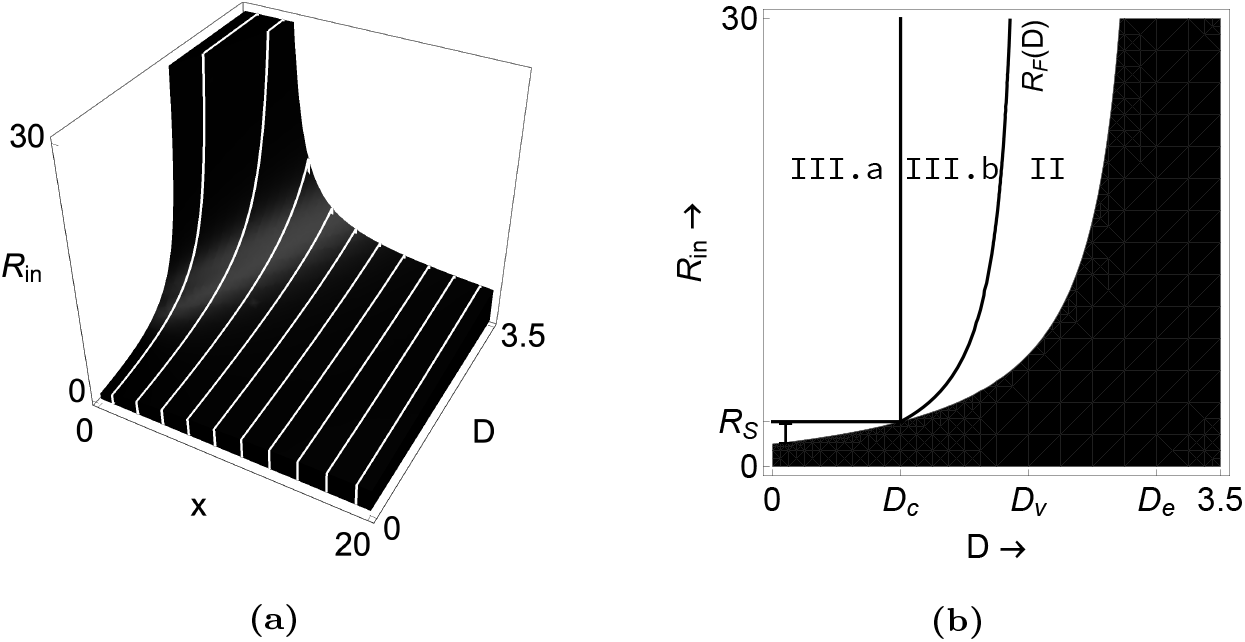
**(a)** Positivity (the empty region) of the invasion fitness in the virgin environment, 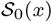, as a function of *D, R_in_* and *x*. **(b)** The cross-section through **(a)** at *x* = 2, in which the empty region with partitions is in order to illustrate the classification I-III.b. In this figure, *μ*_1_ =3, *μ*_2_ = 1.8, *ν*_1_ = *ν*_2_ = 1.5, *δ*_1_ = 1, *δ*_2_ = 1.2, and *K* = 10. Further, in **(b)**, *D_c_* = 1, *D_v_* = 2, *D_e_* = 3, and *R_S_* = 3.

### 3.2. Monomorphic resident dynamics

Suppose that the condition (3) is satisfied so that the population of *x* is viable in the virgin environment. Then the ecological dynamics described by (2) have the following properties:

i. the system is permanent;
ii. a unique positive equilibrium *E*_+_ = (*R*^*^, *F*^*^, *S*^*^) exists, which is globally stable.

The proofs are rather algebraic and are found in Appendices B–D. What is important is the relationship between the two break-even concentrations and the nutrient concentration in equilibrium:

a. for the *uB-/uaB*-settling,

*R_F_* < *R*^*^ < *R_S_* for case I,
*R_S_* < *R*^*^ < *R_F_* for case II,
*R_u_* < *R*^*^ with *R_F_* < *R_u_* < *R_S_* for case III.a,
*R_u_* < *R*^*^ with *R_S_* < *R_u_* < *R_F_* for case III.b;
b. for the *b*-settling,

*R_F_* < *R*^*^ < *R_S_* for cases I, III.a,
*R_S_* < *R*^*^ < *R_F_* for cases II, III.b.

The classification of nutrient concentration *R*^*^ in equilibrium intuitively determines the sign of the selection gradient for different scenarios, which further dominates the evolutionary outcomes as we will see in Sect. 4.2.

## 4. Evolution of settling rate *x*

In the previous section, we study the ecological dynamics of a single microbe type with different settling mechanisms. We find that no qualitative changes in the monomorphic resident population dynamics, no matter how the settling mechanisms and physical operating conditions change. However, this analysis fails to provide any information on whether the evolutionary dynamics are also insensitive to changes in the settling mechanisms and physical operating conditions. We now study the evolution of the settling rate with different settling mechanisms and investigate how the physical operating conditions affect the course of evolution.

From (1) and (2), the dynamics of an initially rare mutant (*F_m_, S_m_*) with settling rate *y* in environment *e* generated by the monomorphic resident *x* are given by the following linear differential equations:

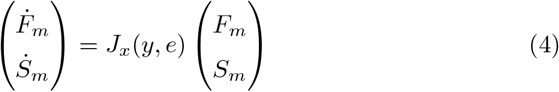

with

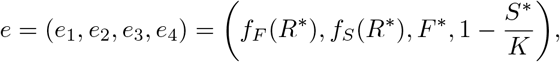

where *e*_1_ is the concentration of nutrient uptake by a floating mutant individual, *e*_2_ is the concentration of nutrient uptake by a settled mutant individual, *e*_3_ is the biomass concentration of resident floaters, *e*_4_ is the vacancy rate of sites on the wall of the chemostat, and *J_x_*(*y, e*) is the per-capita population growth rate of mutant *y* in resident environment (*x, e*). Here we should highlight is that different settling mechanisms have different forms of *J_x_*(*y, e*). More precisely,

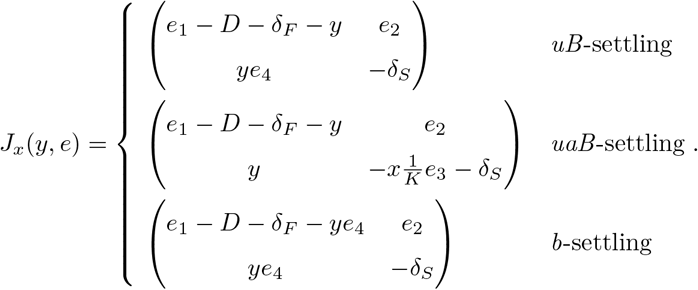

Notice that (4) holds as long as the mutant population remains rare.

### 4.1. Mutant’s fitness

The invasion fitness proxy of mutant *y* in resident *x*, denoted by 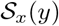, that is

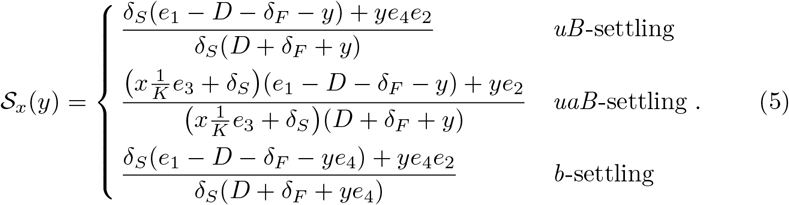

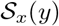 is obtained by calculating – 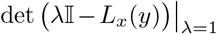 where *L_x_*(*y*) is the next-generation matrix of the population dynamics of mutant *y* in resident environment (*x, e*). Due to different *J_x_*(*y*)(*e*) in different settling mechanisms, the corresponding next-generation matrix *L_x_*(*y*) is different. Consequently, 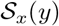 has different forms for different scenarios (ref to Appendix E). As shown in Metz and Leimar (2011) and Appendix E, the mutant can invade the resident if 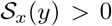, otherwise can not invade if 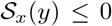. If *y* = *x*, then we have 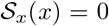 which is so-called *the principle of selective neutrality of residents* in the framework of adaptive dynamics.

From (5), multiple aspects of the resident environment influence the invasion fitness. It implies that the population may become dimorphism, i.e., the mutant *y* coexists with the resident *x* provided that they can mutual invade (Geritz 2005; Dercole and Geritz 2016; Cai and Geritz 2019). However, we find that evolutionary coexistence is impossible in such a chemostat model with dynamics described by (1) as the following section shows.

### 4.2. Evolutionary analysis: direction of evolution, transformation of PIP-types

The direction of gradual evolution is determined by the *selection gradient* at strategy *x*, denoted by 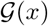, that is

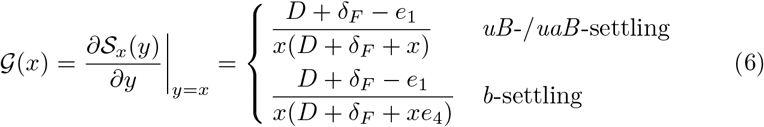

(ref to Appendix F). Notably, the expression 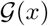 is the same for cases *uB*-settling and *uaB*-settling. The directional evolution proceeding by invasion and substitution leads to the settling rate become a larger value than *x* if 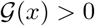. On the contrary, the settling rate evolves to a smaller value than *x* if 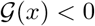. A strategy *x*^*^ is an *evolutionarily singular strategy* if

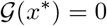

(ref to Geritz et al. 1998). To investigate the evolutionary stability and convergence stability of evolutionarily singular strategies in the strategy space, we have to look at the second-order derivative of (5). From Geritz et al. (1998), if a evolutionarily singular strategy *x*^*^ is a (local) fitness maximum, it is a (local) *evolutionarily stable strategy* (ESS) satisfying

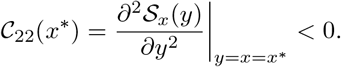

The evolutionarily singular strategy *x*^*^ is *convergence stable* if

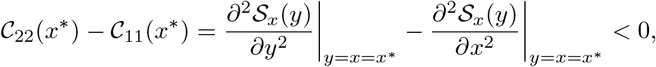

which is an attractor of the strategy substitution process. Conversely, if *x*^*^ is not convergence stable, it is an *evolutionary repeller* (ER). If an evolutionarily singular strategy is both evolutionarily stable and convergence stable, it is a *continuously stable strategy* (CSS). When the gradual evolution process towards an evolutionarily singular strategy that is not evolutionarily stable but convergence stable, it implies that any mutant nearby the evolutionarily singular strategy can invade. Therefore, such an evolutionarily singular strategy is an *evolutionary branching point* (BP) in the strategy space.

From (6), the sign of 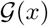 is completely determined by the term *D* + *δ_F_* – *e*_1_. By the classification of the nutrient concentration in the resident equilibrium (see (*a*) and (*b*) in Sect. 3.2), we conclude the signs of 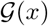 for different scenarios in Tab. 2.

**Tab. 2.**
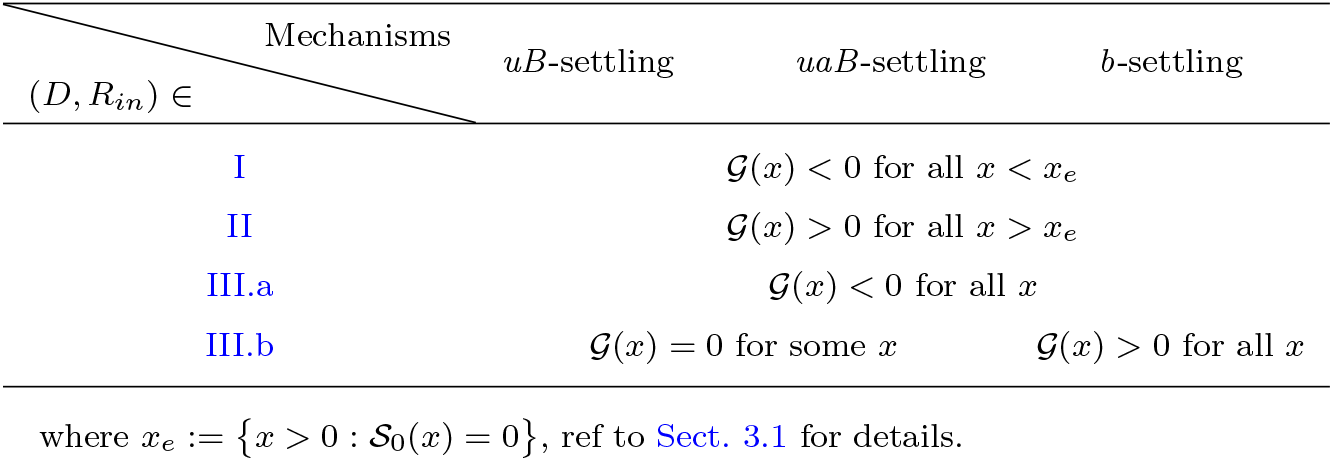
Signs of selection gradient 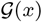 for different scenarios.

If the physical operating conditions (*D, R_in_*) satisfy case I, 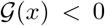 for all *x* < *x_e_* and for all three settling mechanisms, which implies that a mutant with a smaller value of settling rate always can invade and eventually substitute the resident. Thus, the long-term evolution leads to the settling rate approach zero, which means that the whole population finally float in the fluid of the chemostat. Conversely, in case II, 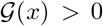 for all *x* > *x_e_* and for all three settling mechanisms, which implies a mutant with a larger value of settling rate always can invade and eventually substitute the resident. Therefore, the longterm evolution leads to the settling rate approach infinity, which means that the whole population finally settle down on the wall of the chemostat. For cases III.a and III.b, the dilution operations and settling mechanisms significantly affect the course of evolution. More precisely, in case III.a, the selection gradient still keeps the same sign whatever the settling mechanism is. The long-term evolution leads to the settling rate reach zero because of 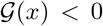 for all *x*. However, the settling mechanisms influence the evolutionary dynamics if an over-dilution operation is performed such that case III.b is satisfied. (*i*) For the *uB-/uaB*-settling, by the nutrient concentration in the monomorphic resident equilibrium (i.e., *R_u_* < *R*^*^ with *R_S_* < *R_u_* < *R_F_*) and (6), we see that 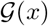 may vanish for some *x*. In other words, evolutionary singular strategies may appear. Notice that the numerator of 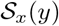 is linear in *y*, and direct calculations give 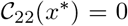 for all possible evolutionary singular strategies *x*^*^. It implies that the evolutionary branching of the strategy substitution process will not occur. Therefore, evolutionary coexistence is impossible. (ii) For the *b*-settling, 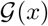 is always positive, which implies that the long-term evolution leads to the settling rate approach infinity.

From the above analysis, evolutionarily singular strategies only occur in the uB-settling and *uaB*-settling and when (*D, R_in_*) satisfy III.b. Fig. 3a shows the number and stability properties of evolutionarily singular strategies in terms of *D* and *R_in_*. Evolutionarily singular strategies, *x*^*^, only exist in the blue region, which are all single (i.e., linear in (*D, R_in_*)) and CSSs. Fig. 3b gives a cross-section through Fig. 3a at *R_in_* = 12 to show how the actual values of *x*^*^ depend on *D*. The values of *x*^*^ gradually increase if the dilution rate increases and then approach infinity if the dilution rate is close to 1.61. Fig. 3c gives a cross-section through Fig. 3a at *D* = 1.2 ∈ (*D_c_, D_v_*). The values of *x*^*^ gradually decrease from infinity to zero if the concentration of nutrient input continuously increases from 5.1.

**Fig. 3.**
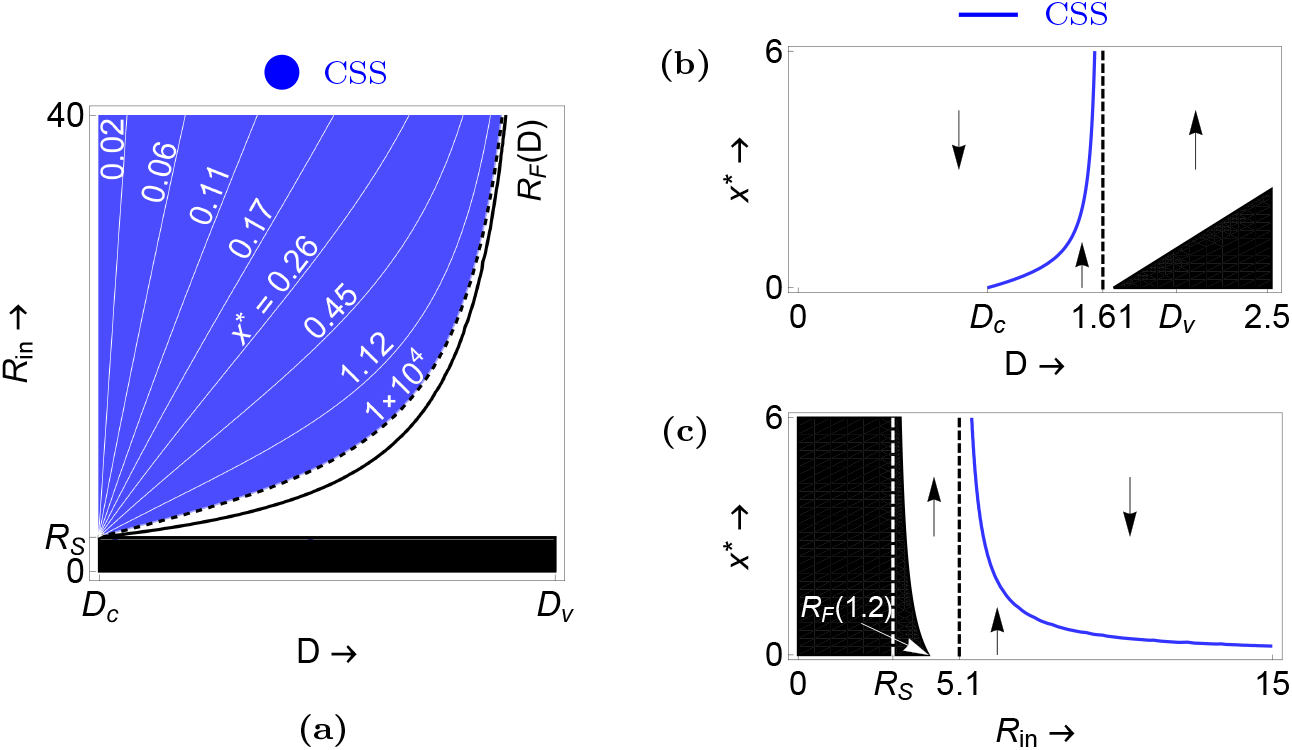
(Colour on the web version) **(a)** Number and stability properties of the evolutionarily singular strategies as a function of *D* and *R_in_* for the *uB*-/*uaB*-settling. In the blue region: the solid lines are contours of the evolutionarily singular strategies, and the dashed line on the boundary corresponds to the sufficiently-high-valued contour *x*^*^ = 1 × 10^4^. **(b)** The actual values of the evolutionarily singular strategies for fixed *R_in_* = 12, and **(c)** *D* = 1.2. In **(b)** and **(c)**, the arrows obtained from the monomorphic selection gradient (6) indicate the evolutionary directions in a monomorphic environment, and the dashed black lines correspond to the points on the dashed line of **(a)**. CSS = continuously stable strategy. The other parameters are the same as Fig. 2.

Fig. 4 concludes the types of Pairwise Invasibility Plots (PIP) for different settling mechanisms and physical operating conditions. For the *uB*-/*uaB*-settling, the PIP-types gradually change (i.e., the vertical invasion boundary through *x*^*^ moves gradually) if the dilution operation moves from an under-dilution to an over-dilution and the dilution rate continuously increases, we refer this transformation to as “gradual transformation” (see Fig. 4a). For the *b*-settling, the PIP-types suddenly change if the dilution operation switches from an under-dilution to an over-dilution (or the other way around), we refer this transformation to as “bang-bang transformation” (see Fig. 4b).

**Fig. 4.**
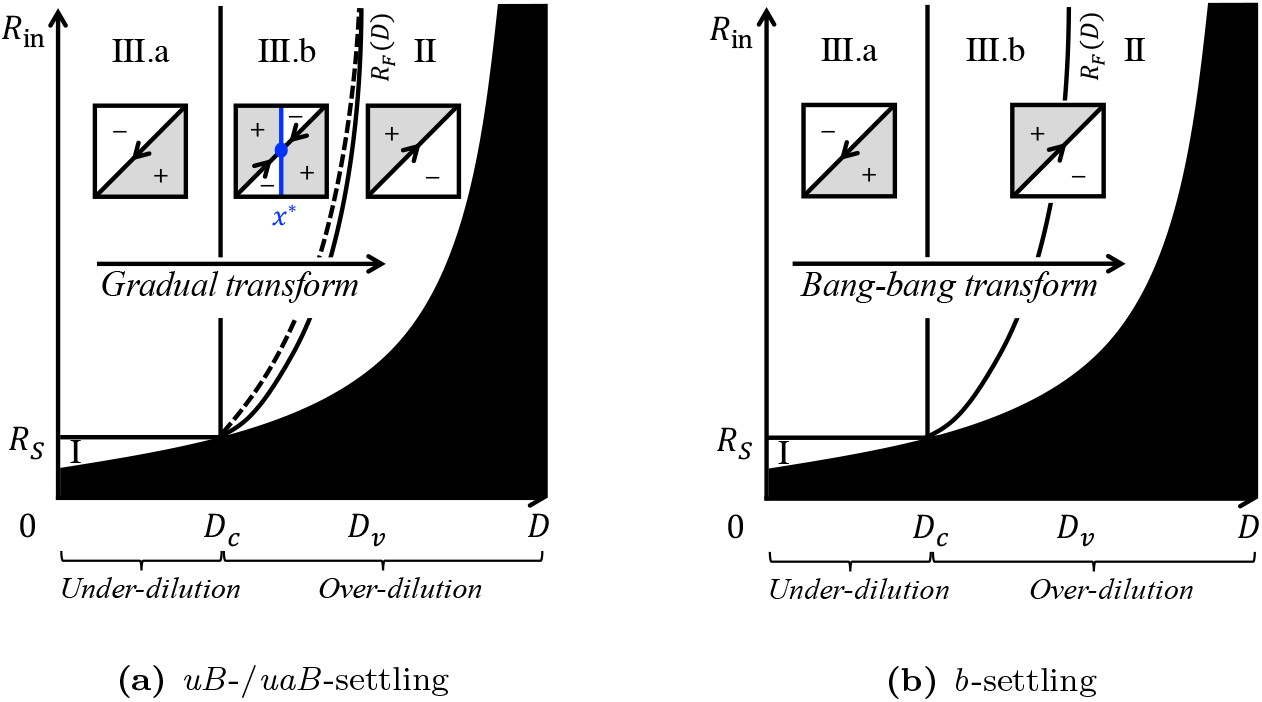
The pairwise invasibility plots (PIP) and the transformation of PIP-types for different scenarios. In each PIP shaded regions depict the mutant strategies that can invade the resident strategy. See text for further explanation.

## 5. Evolution coexistence in a fluctuating environment

As the previous section has shown, the microbial population is always monomorphic during the long-term evolution, whatever the settling mechanism and physical operating condition are. Our interest is whether the microbial population with a particular settling mechanism can become dimorphism or even polymorphism in a fluctuating environment. Especially, how does the microbial community coexist if it is polymorphism? We now investigate the evolution of the settling rate in a fluctuating environment.

To construct a fluctuating environment, one way is to allow the nutrient input to periodically vary for simulating seasons or day/night cycles in ecology. We thus define a sinusoidal perturbation of the nutrient input by

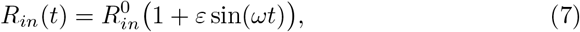

where 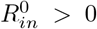 is the average value of *R_in_*(*t*), *ε* ∈ [0,1] is the “degree” of periodicity, and *ω* > 0 is the frequency (Kuznetsov et al. 1992). Notice that 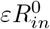 is the amplitude of the periodic perturbation, and 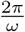 (denote as *T*) is the period.

In such a fluctuating environment, the ecological dynamics of a monomorphic population in the chemostat is still described by (2) but with periodic nutrient input (7). To avoid redundancy, we do not repeat these differential equations here. Nevertheless, the definitions of one steady-state and two matrices have to highlight throughout this section. (*i*) The washout steady-state 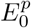 (with respect to *E*_0_ in Sect. 3.1) is the periodic solution of differential equations describing the physical dynamics of nutrient with periodic input (7). (*ii*) The parameter matrix 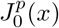 (with respect to *J*_0_(*x*) in (A) of Appendix A) is the per-capita growth rate matrix of the population of *x* in the virgin environment that is generated by the physical dynamics of nutrient with periodic input (7). (*iii*) The parameter matrix 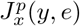 (with respect to *J_x_*(*y, e*) in (4)) is the percapita growth rate matrix of mutant *y* in resident environment (*x, e*) that is generated by resident x in the fluctuating environment. The following we apply the theory of Floquet to the analysis of the evolutionary dynamics.

### 5.1. Monomorphic resident dynamics

In the virgin environment, an initially rare microbial population with settling rate *x* is viable if

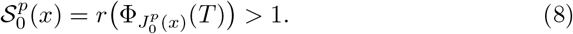

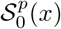 is defined by the maximum modulus of characteristic (or Floquet) multipliers of the corresponding periodically forced system, which is the maximum modulus of eigenvalues of the monodromy matrix 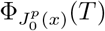. Here 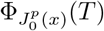 is obtained by integration over one period of the variational equations around 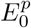 with initial conditions the unit matrix (see, e.g., Lenas and Pavlou 1995; Klausmeier 2008).

Once the microbial population of x is viable in the virgin environment, the periodically forced system admits at least a positive *T*-periodic solution which is uniformly persistent (ref to Appendix G). Notice that, by a standard argument, the solutions of the periodically forced system are also globally bounded. Although we are not able to theoretically prove the uniqueness and global stability of the positive *T*-periodic solution under the condition (8), there exists a *T*-periodic solution that is with respect to the unique positive equilibrium of the unforced system (2).

In Sect. 5.3 and Sect. 5.4, we numerically found a limit cycle of resident x by using the *Mathematica*^®^ software as Lehtinen and Geritz (2019) did. More precisely, fixed the initial value sufficiently close to but not identical to the unique positive equilibrium of the unforced system, we numerically integrated the corresponding differential equations and collected data at each point in time along the orbit. The convergence of the orbit was evaluated using a Poincaré section, which we implemented using the *EventLocater* method for *NDSolve*. The data were collected until the distance between two consecutive equilibrium points of a Poincaré map was smaller than 10^−6^, after which we discarded the transient data. The desired limit cycle was described by the data from the last iteration.

### 5.2. Mutant’s fitness

After successful establishment in the virgin environment, the invasion fitness proxy of an initially rare mutant *y* in the environment generated by resident *x* is defined by

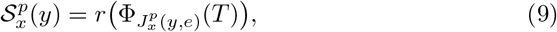

which is the maximum modulus of characteristic (or Floquet) multipliers of the periodically forced system, and 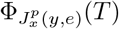 is the monodromy matrix resulting from integration over one period of the variational equations around a resident limit cycle with initial conditions the unit matrix. By the theory of Floquet, we claim that a mutant *y* can invade the resident *x* if 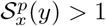, otherwise can not invade if 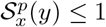.

### 5.3. Number and stability properties of evolutionarily singular strategies

In this section, we investigate how the settling mechanisms and physical operating conditions affect the singular strategies of the evolution with nonequilibrium resident dynamics. Here we only focus on the dilution rate *D*, the averaged concentration of nutrient input 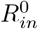, and the frequency of periodic input *ω*.

Fig. 5 gives the number and stability properties of evolutionarily singular strategies in terms of *D*, 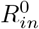 and *ω*. Each subgraph is generated by the projection of rescaled *x*^*^ in the 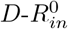 plane for a particular settling mechanism and for fixed *ω*, where the *x*^*^-axis is rescaled by 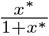 in order to consider all possible strategy values. Fig. 6 gives the cross-sections through a particular subgraph of Fig. 5 at fixed 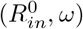 to help interpret how the actual values of the rescaled evolutionarily singular strategy changes in terms of *D*. As we see, each vertical axis in Fig. 6 is the linear transformation of *x*\* used in Fig. 5, i.e., 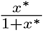. Next, we specify each subgraph of Figs. 5 and 6.

**Fig. 5.**
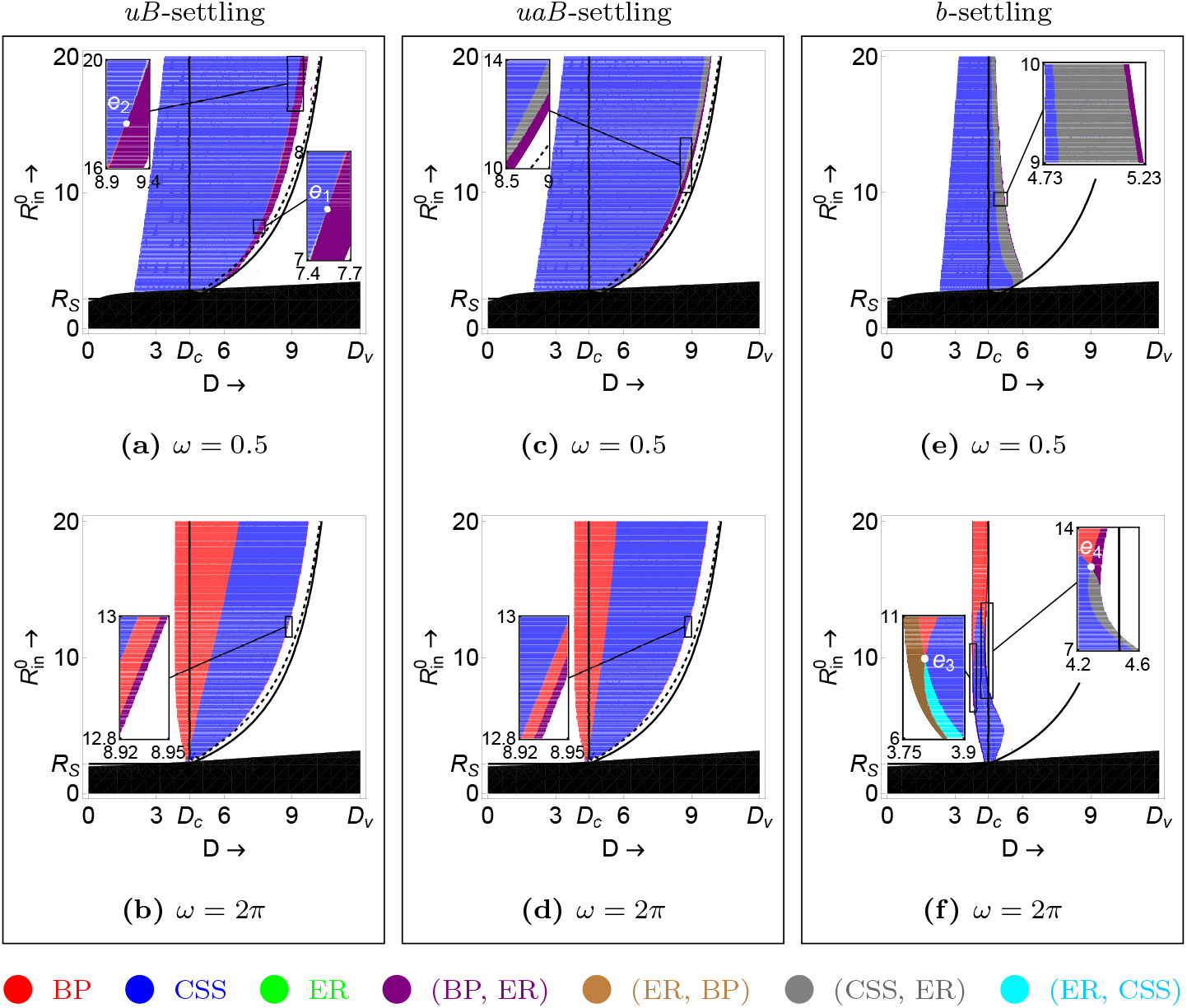
(Colour on the web version) Number and stability properties of the evolutionarily singular strategies as a function of *D* and 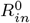 for a particular settling mechanism in low-frequency (*ω* = 0.5) and high-frequency (*ω* = 2π) inputs, respectively. In each plot the solid lines and the dashed line correspond to that in Fig. 4. BP = evolutionary branching point, CSS = continuously stable strategy, ER = evolutionary repeller, and an ordered combination of two of them denotes double singular strategies such as (BP, ER) which indicates that the small-valued singularity is a BP and the large-valued singularity is an ER. Notice different colour regions correspond to different numbers and stability properties of the evolutionarily singular strategies. See text for further explanation. In this figure, *μ*_1_ = 13, *μ*_2_ = 2.3, *ν*_1_ = 3, *ν*_2_ = 2, *δ*_1_ = 1, *δ*_2_ = 1.2, *K* = 10, and *ε* = 0.9. Further, *D_c_* = 4.47368, *D_v_* = 12, and *R_S_* = 2.18182. The intersection points *e*_1_ = (7.54, 7.47), *e*_2_ = (9.18, 17.65), *e*_3_ = (3.8, 9.27), and *e*_4_ = (4.28, 11.78).

**Fig. 6.**
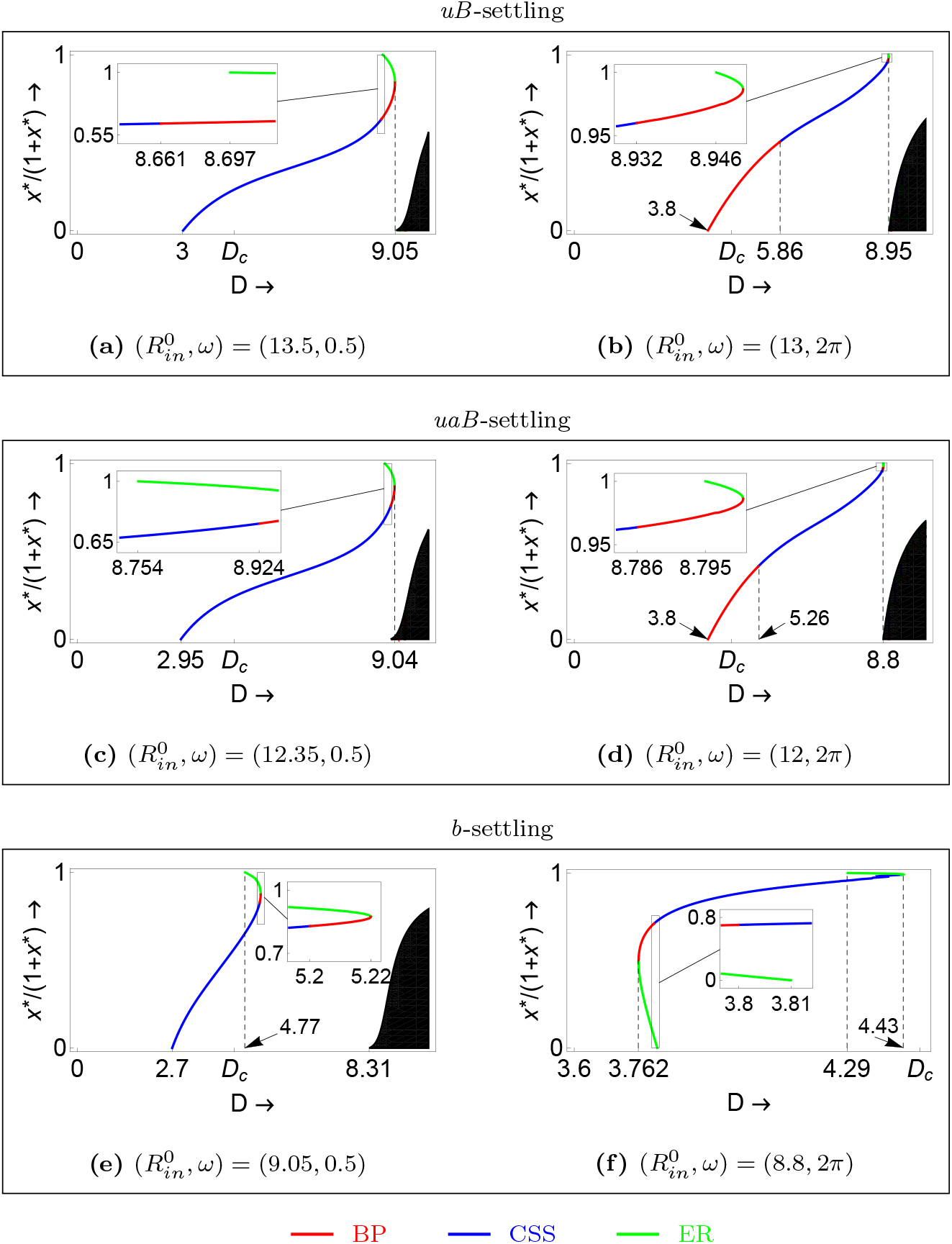
(Colour on the web version) The actual values of a linear transformation of the evolutionarily singular strategy as a function of *D* for fixed 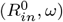 and a particular settling mechanism. See text for further explanation. The other parameters are the same as Fig. 5.

i. For the case of *uB*-settling: Fig. 5a shows that the singular strategies present different stability properties in a low-frequency (*ω* = 0.5) input. Recall the evolutionary dynamics in the environment with constant input (ref to Sect. 4.2), the singular strategy occurs only when we perform the over-dilution operation with 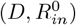 satisfying case III.b. In the low-frequency input, however, the singular strategy may occur and is always a CSS when we perform the under-dilution operation with 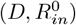 satisfying case III.a. Double singular strategies appear when 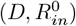 are close to the dashed line. Fig. 6a gives the cross-section at 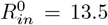, which shows the singular strategy is a CSS if *D* ∈ [3, 8.661) and then is an EP if *D* ∈ [8.661, 8.697). Further, the double singular strategies appear if *D* ∈ [8.697, 9.05], which is a combination of a small-valued BP and a large-valued ER. Once *D* is larger than 9.05, the two singular strategies vanishes. This type of bifurcation of singular strategies only happens for 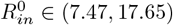 (ref to the two intersection points in Fig. 5a). Consider a cross-section at a fixed 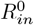 with value either smaller than 7.47 or larger than 17.65, the type of double singular strategies may become a different combination that consists of a small-valued CSS and a large-valued ER. In a high-frequency (*ω* = 2*π*) input, Fig. 5b shows a significant difference. BPs always occur when we perform a dilution operation with values around the critical rate *D_c_*. Fig. 6b is the cross-section at 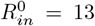. The singular strategy is a BP if *D* ∈ [3.8, 5.86), and becomes a CSS if *D* ∈ [5.86, 8.932), but returns to a BP if *D* ∈ [8.932, 8.946). Continuously increasing *D*, double singular strategies occur for *D* ∈ [8.946, 8.95] which is a combination of a small-valued BP and a large-valued ER. The two singular strategies disappear as soon as *D* is larger than 8.95.
ii. For the case of *uaB*-settling: In the low-frequency input with 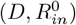 close to the dashed line as Fig. 5c shows, the situation is different with that of the *uB*-settling. Fig. 6c gives the cross-section at 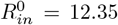, which shows the singular strategy exists for *D* ∈ [2.95, 9.04]. The singular strategy is a CSS if *D* ∈ [2.95, 8.754). The number of singular strategies becomes twice if *D* ∈ [8.754, 9.04]. Meanwhile, we find that the combination of double singular strategies may change for different dilution rates. More precisely, it is consist of a small-valued CSS and a large-valued ER for *D* ∈ [8.754, 8.924). However, if *D* ∈ [8.924, 9.04], the bottom one becomes a BP and the top one is still an ER. In the high-frequency input, the bifurcation of singular strategies is similar to the case of *uB*-settling as Fig. 5d shows. The unique difference is the parameter region for different stability properties of the singular strategies.
iii. For the case of *b*-settling: In the low-frequency input, the singular strategies occur under some parameter conditions, which never happen for the evolution in the environment with constant input (ref to Sect. 4.2). Compare to the *uB*-settling and *uaB*-settling, the difference in Fig. 5e is that singular strategies exist only when the dilution rate is close to the critical rate *D_c_* (N.B., this situation also happens in the high-frequency input). Fig. 6e gives the cross-section at 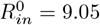, which shows that the singular strategy exists if *D* ∈ [2.7, 5.22] and is a CSS if *D* is less than 4.77, but double singular strategies appear if *D* is larger than 4. 77. For the double singular strategy, it is a combination of a small-valued CSS and a largevalued ER as *D* ∈ [4.77, 5.2) but becomes a different combination that consists of a small-valued BP and a large-valued ER as *D* ∈ [5.2, 5.22]. In the high-frequency input as Fig. 5f shows, it presents a complex bifurcation of singular strategies. The number of singular strategies can change from two to one and then return to two. Fig. 6f gives the cross-section at 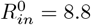, which shows an example of the bifurcation of singular strategies in the number and also in the stability properties. More precisely, the singular strategies exist only for *D* ∈ [3.762,4.43]. Double singular strategies appear if *D* takes value in either [3.762, 3.81] or [4.29,4.43], and it degenerates to the single singular strategy if *D* ∈ (3.81, 4.29). For *D* increases from 3.762, the double singular strategies are consist of a small-valued ER and a large-valued BP. When *D* is larger than 3.8 but less than 3.81, the large-valued singular strategy becomes a CSS. But when *D* is larger than 3.81, the small-valued singular strategy vanishes, and thus only the large-valued CSS exists. The number of singular strategies remains single if *D* continuously increases from 3.81 to 4.29. Once *D* is beyond 4.29, double singular strategies occurs again. Nevertheless, it is a combination of the original CSS and a larger-valued ER. Fig. 6f is just one type of bifurcation of singular strategies from Fig. 5f. One can find more exciting bifurcations of singular strategies in the number and stability properties by selecting different cross-sections through Fig. 5f.

From the above numerical analysis, we see that evolutionarily singular strategies present diversified combinations in the number and stability properties. Notably, different from the constant environment, the long-term evolution for all three settling mechanisms in a fluctuating environment can produce evolutionary branching in the strategy space.

### 5.4. Evolutionary coexistence and the way of coexistence

In the previous section, we see that suitable physical operating conditions will lead to evolutionary branching in the phenotype of the settling rate. However, the evolutionary outcomes after first branching are still unknown. This section is to investigate whether two distinct types of species can stabilise coexist in the long-term evolution and what phenotypes they are.

Suppose that the monomorphic evolution has reached an evolutionary branching point, the gradual evolution further induces a dimorphism of two different strategies, say *x*_1_ and *x*_2_. The study of dimorphic evolution is based on the invasion fitness proxy of a mutant *y* in the dimorphic resident *x*_1_ and *x*_2_:

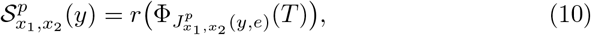

which is the maximum modulus of characteristic (or Floquet) multipliers of the periodically forced system, and 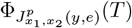 is the monodromy matrix resulting from integration over one period of the variational equations around the dimorphic resident limit cycle with initial conditions the unit matrix. Thus, the mutant *y* can invade either *x*_1_ or *x*_2_ if 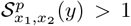, otherwise can not invade if 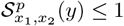. The directions of evolution of resident strategies *x*_1_ and *x*_2_ are determined by the selection gradient

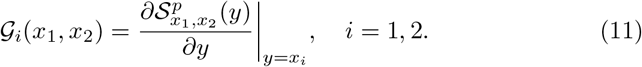

Fig. 7 shows examples of the evolutionary dynamics after first branching for different settling mechanisms and different physical operating conditions. In the first column, corresponding to examples in the low-frequency input, the shaded regions indicate that *x*_1_ and *x*_2_ can mutually invade and hence can coexist as a protected dimorphism. The arrows on the main diagonal show the directions of evolution in the monomorphic resident environment. Once the evolution of the settling rate reaches a BP 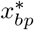, it undergoes disruptive coevolution with the population contains two strategies. Inside the region of coexistence, the evolution in a dimorphic resident environment follows the arrows. Since the evolutionary isoclines inside the region of coexistence do not intersect, there are no dimorphic singular strategies. The population evolves along with the arrows until it reaches the boundary of the coexistence region. On the boundary, the black points indicate the endpoints of dimorphic evolution. The second column is the corresponding fitness at each endpoint, which implies that all endpoints cannot be invaded by any other neighbouring strategies. Consequently, (*x*^**^, ∞) is the evolutionary endpoint. In words, the long-term evolution in the environment with low-frequency input leads to dimorphic coexistence for all three settling mechanisms, in which one of the two coexisting species has to settle down totally on the wall of the chemostat. The third and fourth columns are dimorphic evolution in the environment with the high-frequency input. The difference here is that the evolutionary endpoints all become (0, *x*^**^). That is to say, one of the two coexisting species has to float entirely in the fluid of the chemostat.

**Fig. 7.**
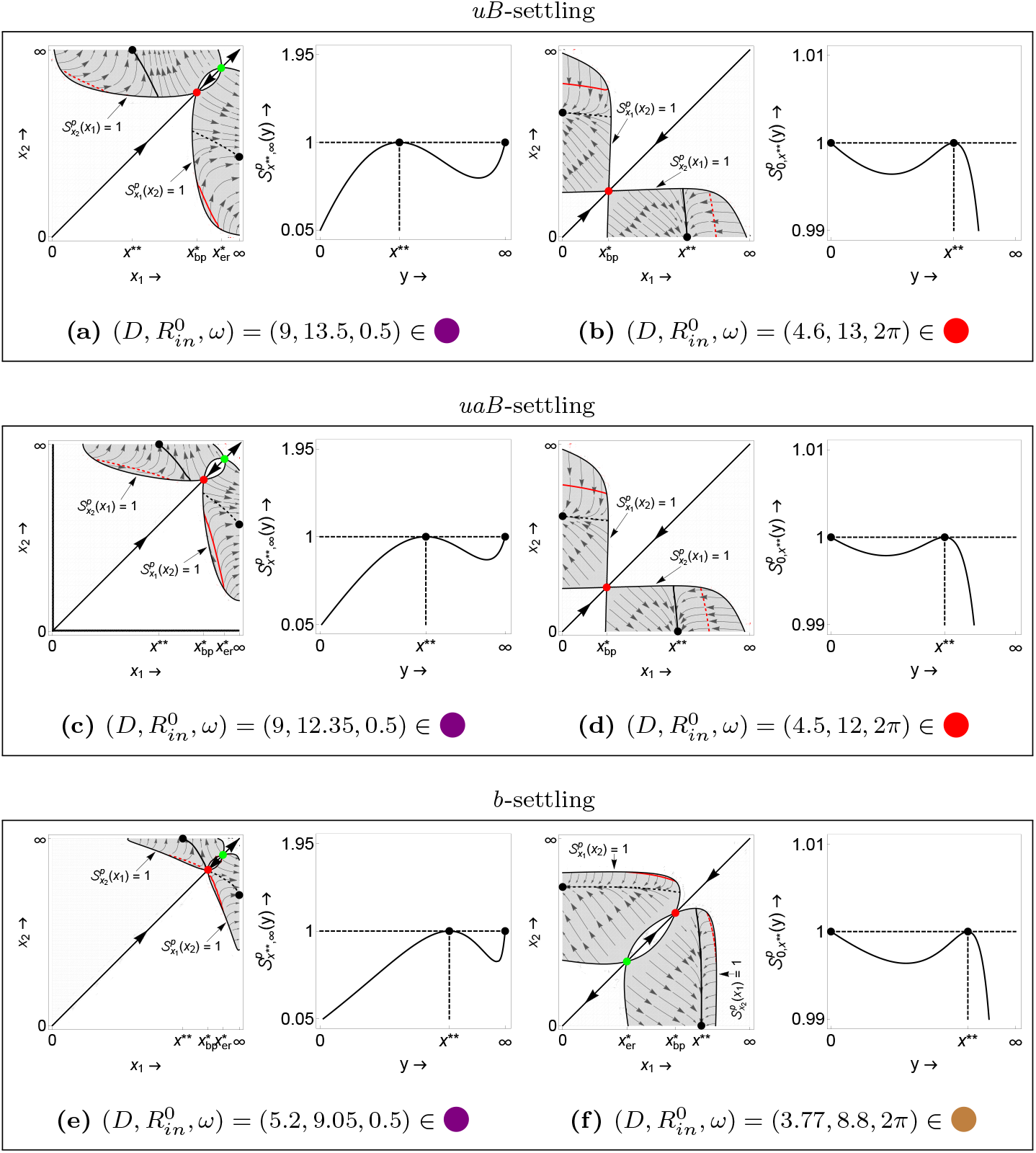
(Colour on the web version) Examples of mutual invasibility plots (MIP) and fitness landscape at the evolutionary endpoint (black dot) for different scenarios. In each MIP: the arrows on the main diagonal show the evolutionary directions in a monomorphic environment; the green dot is an evolutionary repeller and the red dot is an evolutionary branching point; the shaded regions indicating protected dimorphism are separated by stable (black) and unstable (red) isoclines at which selection gradient vanishes in either the *x*_1_-direction (solid) or the *x*_2_-direction (dashed); the vector fields obtained from (11) denote the directions of evolutionary dynamics of strategies *x*_1_ and *x*_2_. Notice that the axes *x*_1_, *x*_2_ and *y* are all rescaled by the same linear transformation as in Fig. 6 in order to consider all strategy values. 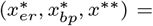 **(a)** (8.56, 3.302, 0.755), **(b)** (–, 0.335,1.958), **(c)** (10.473, 4.107, 1.326), **(d)** (–, 0.317, 1.588), **(e)** (9.49, 4.757, 2.272), **(f)** (0.53, 1.504, 2.797). The other parameters are the same as Fig. 5.

## 6. Conclusion

Site choice and competition (for breeding, foraging, refuge, etc) is an essential component in the lives of many organisms, which is why investigating the mechanism is important. To understand the choice and competition of sites in nature, we consider a microbial population who can settle down on the wall of the chemostat through one of the three settling mechanisms. Employing the framework of adaptive dynamics, we have explored the evolutionary dynamics of the settling rate under different settling mechanisms, respectively. Meanwhile, we also have investigated how physical operating conditions affect the course of evolution. Next, we conclude our main results.

First, we study the ecological dynamics with a single microbe type in the constant environment. We have shown that the classification of nutrient concentration in resident equilibrium intuitively determines the direction of evolution, which further dominates the evolutionary outcomes. The theoretical analysis shows that evolutionary coexistence is impossible in the constant environment for all three settling mechanisms. The long-term evolution induces that the monomorphic population eventually possess zero-, or finite-, or infinity-valued settling rate, which depends on the settling mechanisms and physical operating conditions. In other words, the composition of the population is dominated by the settling mechanisms and physical operating conditions through the evolutionary dynamics. Fig. 4 concludes the evolutionary outcomes for different scenarios. In particular, if both floaters and settlers have positive intrinsic growth rates when they exist alone in the virgin environment, the direction of monomorphic evolution will significantly change when the dilution operation varies around the critical dilution rate. More precisely, (*i*) if the population adopts either the *uB*-settling or the *uaB*-settling, the successive evolution causes that the settling rate reaches a continuously stable strategy. Moreover, its values will gradually move from zero to infinity if the dilution rate continuously increases. The types of Pairwise Invaisbility Plots present a gradual transformation. (*ii*) If the population takes a way of *b*-settling, the direction of evolution will suddenly reverse when the dilution operation varies around the critical dilution rate. It corresponds to the bang-bang transformation of types of Pairwise Invaisbility Plots.

Second, we periodically vary the nutrient input to investigate the evolution of the settling rate in a fluctuating environment. The numerical analysis shows that if the dilution rate, the averaged concentration of nutrient input, and the frequency of input are in a proper range, evolutionary branching in the phenotype of the settling rate will occur. It opens the possibility of evolutionary coexistence for two distinct phenotypes. After first evolutionary branching, two phenotypes can eventually evolve into two generalist species for all three settling mechanisms. However, the difference is the way they coexist in different frequency inputs. In a low-frequency input, the long-term evolution leads to one of two coexisting species settle down totally on the wall of the chemostat. Conversely, the long-term evolution in the environment with a high-frequency input causes one of the two coexisting species to float entirely in the fluid of the chemostat. Consequently, the settling mechanisms and physical operating conditions dominate the phenotypic diversity and coexistence way of the microbial population in a fluctuating environment.

In this paper, we have investigated the evolution of the settling rate under three different mechanisms involving site choice and competition. However, in reality, matters are more complicated, especially the population can adopt a mixed settling strategy to respond to changes in the environment. A question that arises here is “what does cost the population has to pay for altering its behaviour”. Furthermore, the evolutionary dynamics are also worth investigating. We leave these as subsequent works.

## Acknowledgements

YC was supported by China Scholarship Council Grant 201406650003.

## Appendix A The invasion fitness proxy of *x* in the virgin environment, 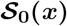

The population dynamics of *x* in the virgin environment (i.e., at the washout equilibrium *E*_0_) is given by

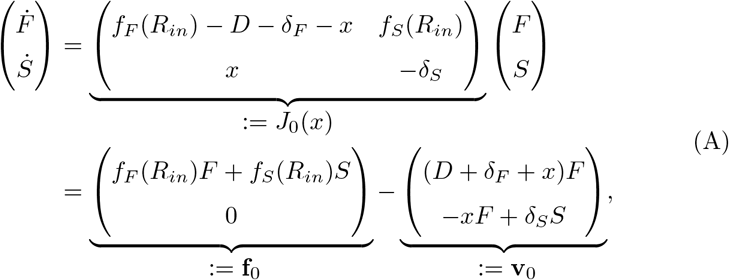

which is the same for all three settling mechanisms. The partial derivatives of **f**_0_ and **v**_0_ with respect to (*F, S*), evaluated at the washout equilibrium *E*_0_, which denote as 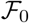 and 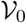 respectively, i.e.,

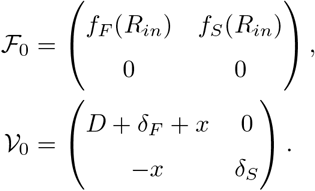

The next-generation matrix *L*_0_(*x*) is defined by 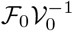 (Diekmann and Heesterbeek 2000), i.e.,

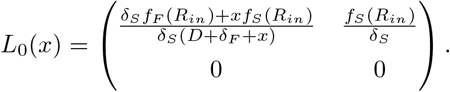

Metz and Leimar (2011) provides an invasion fitness proxy

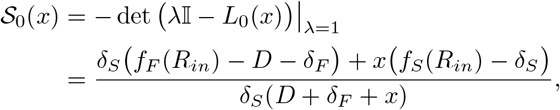

from which the population of *x* is viable if 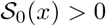 and inviable if 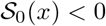.

Since

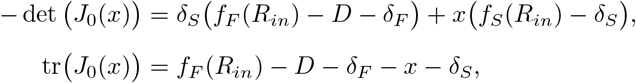

we have

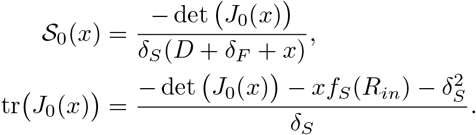

Further, 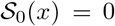 implies – det (*J*_0_(*x*)) = 0 and tr(*J*_0_(*x*)) < 0. Thus, we claim that the population of *x* is viable in the virgin environment if and only if 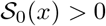.

## Appendix B Permanence of the systems (2)

Let 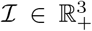, 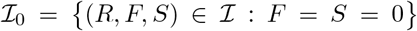 be the extinction set and 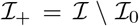 be the space of positive population sizes. The permanence is obtained by the global boundedness and the uniformly persistence of the solutions.

Let function

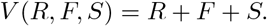

Computing the time derivative of *V* along the solutions 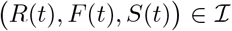, we have

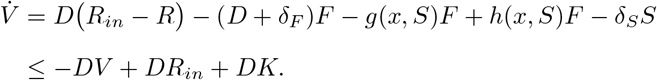

Notice that here we have applied *S*(*t*) ≤ *K* for all *t* ≥ 0. It follows that

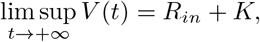

which means the solutions of the system (2) are bounded and well define on 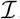. Moreover, it is not difficult to show 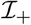 is forward invariant and *R*(*t*) > 0 for all *t* > 0. In words, the solutions (*R*(*t*), *F*(*t*), *S*(*t*)) remain strictly positive for all *t* > 0 if (*R*(0), *F*(0), *S*(0)) is positive.

The uniformly persistence is proved by analyzing the stable and unstable manifolds of *E*_0_ (see, e.g., Hofbauer and So 1989; Lemesle and Gouze 2008; Gouzé and Lemesle 2008). Notice that *E*_0_ is the maximal invariant set in 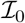. Let 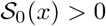, the eigenvalues of the Jacobian matrix evaluating at *E*_0_ satisfy

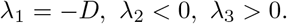

**Fig. B.**
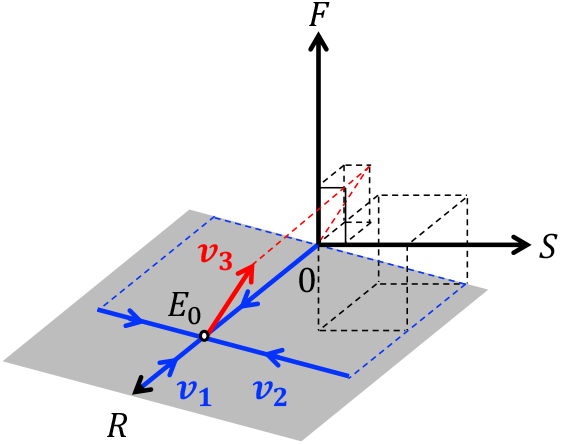
(Colour on the web version) Stable and unstable manifolds of *E*_0_.

The corresponding eigenvectors *v_i_* = (*v*_*i*1_, *v*_*i*2_, *v*_*i*3_)^T^ *i* = 1, 2, 3, are

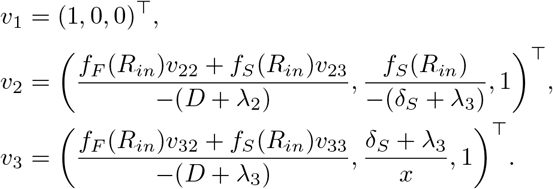

The intersection of the local stable manifold of *E*_0_ and 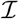 is 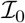, while the local unstable manifold intersects 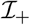 (ref to Fig. B). From Hofbauer and So (1989, Theorem 4.1), 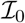 is a repeller. Since there is not heteroclinic cycle in the boundary, we claim that the positive solutions of the system (2) are uniformly persistent, i.e., there exists a positive constant *η* such that

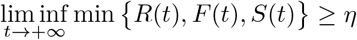

for all initial values 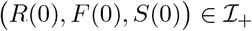.

## Appendix C Existence and uniqueness of the positive equilibrium *E*_+_ of the system (2)

The existence and uniqueness of *E*_+_ is obtained through a geometrical analysis. The basic idea is to reduce the three-dimensional system (2) into the *R*-component. Then it remains to analysis the existence and uniqueness of solutions *R* of the equation 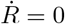.

Solving the equations 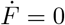 and 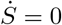, we have where

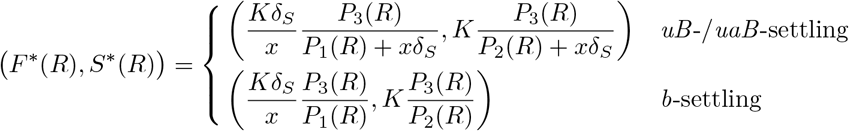

where

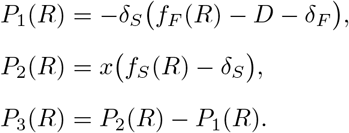

Further, the sufficient and necessary condition for 0 < *F*\*(*R*) and 0 < *S*\*(*R*) < *K* is that there exists *R* > 0,

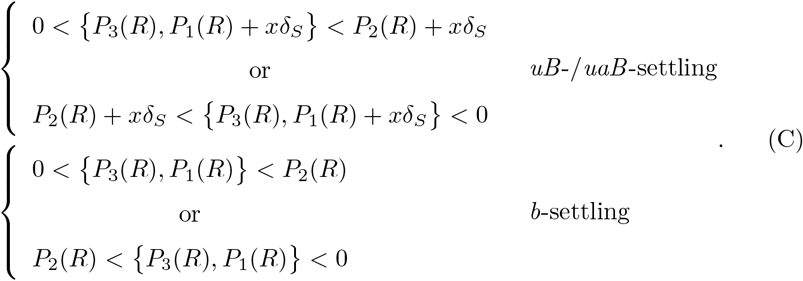

For the case of *uB-/uaB*-settling, if 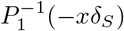 exists, denoted by *R_c_*, then

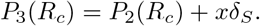

Since *P*_3_(0) < 0 and 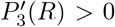 for all *R* > 0, there exists *R_u_* ∈ (0, *R_in_*) such that

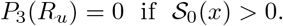

Thus, the first assertion of (C) holds if *R* ∈ (*R_u_*, min{*R_c_, R_in_*}). For the case of *b*-settling, there exists *R_b_* ∈ (0, *R_in_*) such that

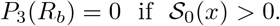

The second assertion of (C) depends on the particular nutrient uptake rates.

The shaded intervals in Fig. C indicate the possible *R* to guarantee *F*^*^(*R*) > 0 and 0 < *S*\*(*R*) < *K* when a particular mechanism is under consideration. Notice *R*(*t*) ≤ *R_in_* for all *t* ≥ 0. We now specify the intervals of *R* that meet different scenarios:

(Ca) for the *uB*-/*uaB*-settling,

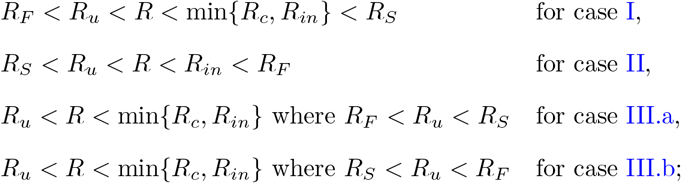
(Cb) for the *b*-settling,

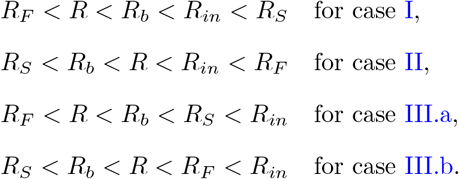

Notice that *R_S_* may not exist in case I (i.e., *R_S_* = +∞), and so is *R_F_* in case II.

Substituting (*F*\*(*R*), *S*\*(*R*)) into the equation 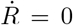, then the existence and uniqueness of *E*_+_ is derived by solutions *R* of the equation 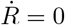. In fact,

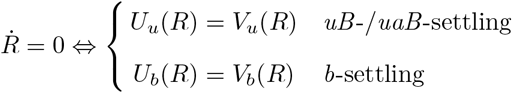

where

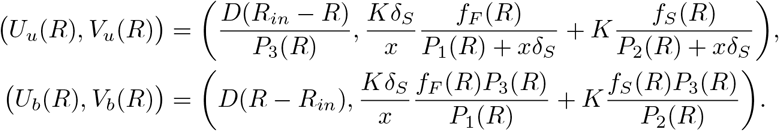

For each interval of *R* in (Ca) and (Cb), it is not difficult to verify the existence and uniqueness of solutions *R* of the equation 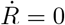 by the monotonicity of *U_u_, V_u_, U_b_* and *V_b_*. Thus, the system (2) admits a unique positive equilibrium *E*_+_ if 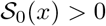.

**Fig. C.**
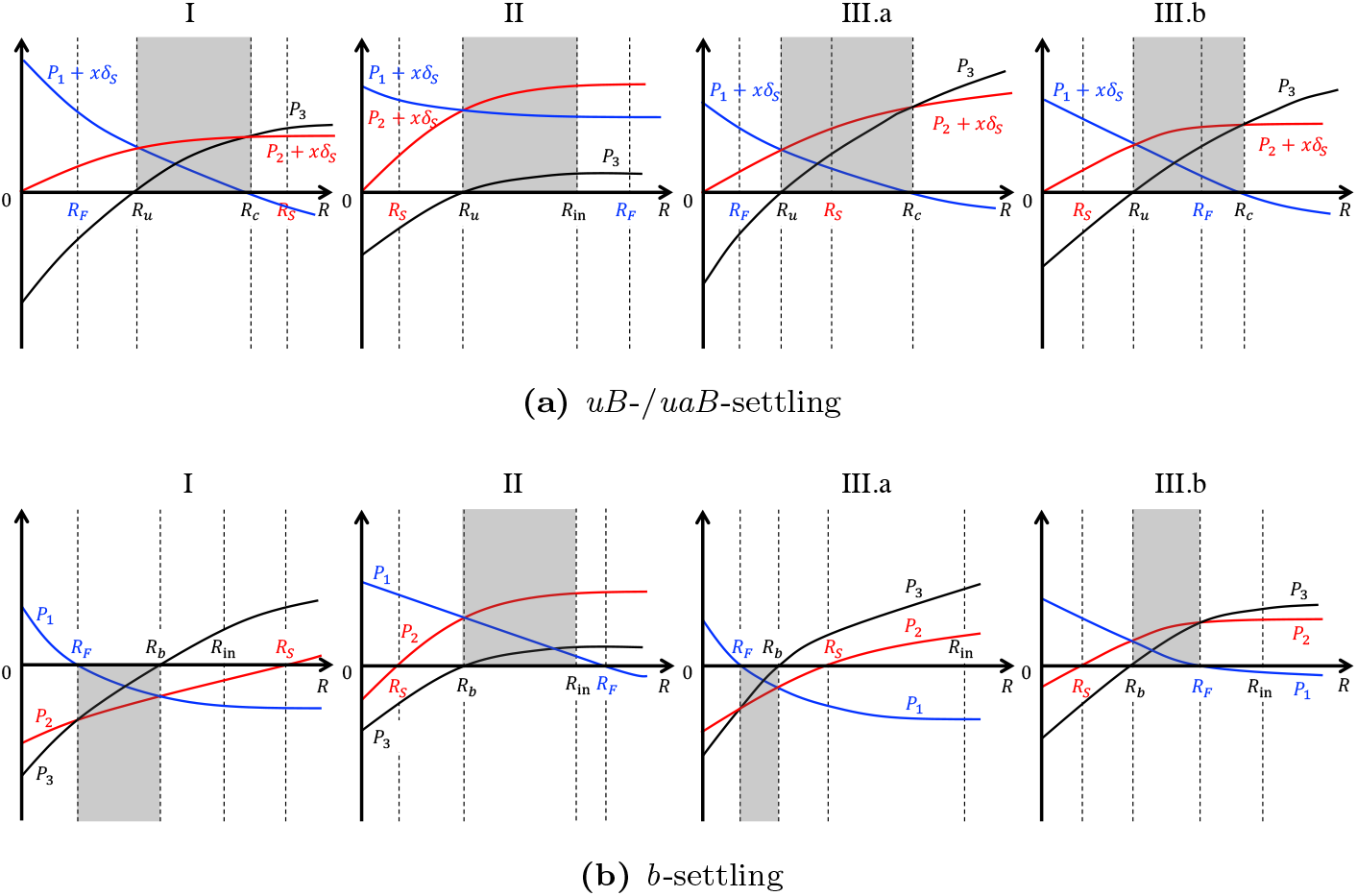
(Colour on the web version) Intervals (shaded regions) of *R* that (C) holds in different scenarios.

## Appendix D Global stability of the positive equilibrium *E*_+_ of the system (2)

The global stability of the positive equilibrium *E*_+_ is obtained by applying the high-dimensional Bendixson’s criterion of Li and Muldowney (1993).

Let 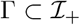 is an open connected set and *n*_0_ = (*R*(0), *F*(0), *S*(0)) ∈ Γ. Let *φ*(*t, n*_0_) is the unique solution of the system (2) with *φ*(0, *n*_0_) = *n*_0_, and *A*^[2]^ is the second additive compound matrix of the corresponding Jacobian matrix *J_x_*(*x*) = (*J_ij_*)_1≤*i,j*≤3_ of the system (2), i.e.,

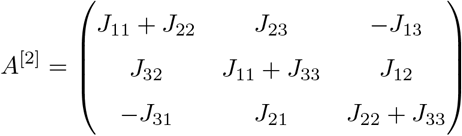

where 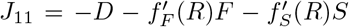, *J*_12_ = – *f_F_*(*R*), *J*_13_ = – *f_S_*(*R*), 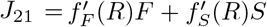, *J*_22_ = *f_F_*(*R*) – *D* – *δ_F_* – *g*(*x, S*), *J*_23_ = *f_S_*(*R*) – *g′*(*x, S*)*F*, *J*_31_ = 0, *J*_32_ = *h*(*x, S*), *J*_33_ = *h′*(*x, S*)*F* – *δ_s_*, and

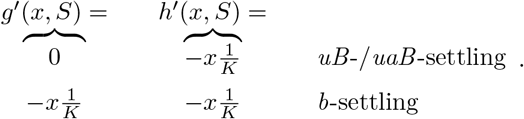

As shown in Li and Muldowney (1993), to derive a high-dimensional Bendixson’s criterion, it is sufficient to show that the second compound equation of the system (2):

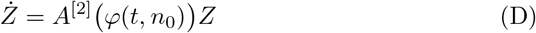

with respect to a solution *φ*(*t, n*_0_) ∈ Γ, which is equi-uniformly asymptotically stable, i.e., for each *n*_0_ ∈ Γ, the system (D) is uniformly asymptotically stable, and the exponential decay rate is uniform for *n*_0_ in each compact subset of Γ. The equil-uniform asymptotic stability of the system (D) implies the exponential decay of the surface area of any compact two-dimensional surface in Γ. If Γ is simply connected, this excludes the existence of any invariant simple closed rectifiable curve in Γ, including periodic orbits, homoclinic orbits, heteroclinic cycles, etc. Moreover, if Γ is positively invariant and contains a unique equilibrium, then the equilibrium is globally asymptotically stable in Γ. To establish the stability, it requires to construct a suitable Lyapunov function (see also Sun and Loreau 2009). More precisely, the system (D) is equi-uniformly asymptotically stable if there exists a positive definite function *V*(*Z*), such that the time derivative 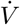 along the solutions of the system (D) is negative definite which is independent of *n*_0_.

Let function

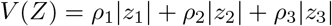

with *ρ*_1_ ≥ *ρ*_2_ ≥ *ρ*_3_ > 0. Calculating the right-hand derivative 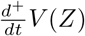, we have

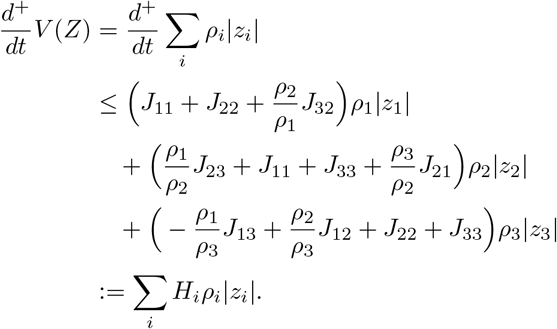

From the permanence of the solutions of the system (2) (see Appendix B), there exist *ξ* > *η* > 0 such that

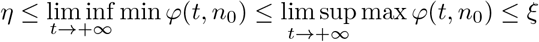

for all 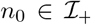. We now fix *ξ* and *η*, and set 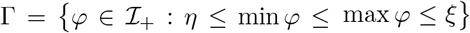. Since *ρ*_1_ ≥ *ρ*_2_ ≥ *ρ*_3_ > 0, we can always find *ρ_i_* for *i* = 1, 2, 3 such that

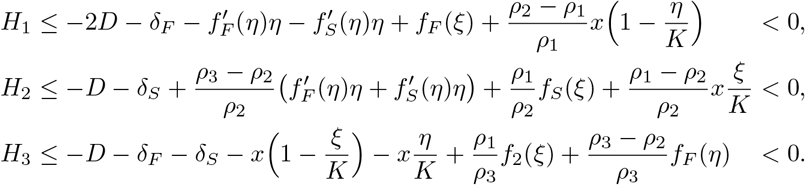

Further,

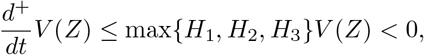

which guarantees the equi-uniform asymptotic stability of the second compound system (D). Hence, by Sun and Loreau (2009, Proposition 4.1), the unique positive equilibrium *E*_+_ of the system (2) is globally stable.

## Appendix E The invasion fitness proxy of a mutant *y* in the resident *x*, 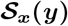

The method to obtain *S_x_*(*y*) is the same as shown in Appendix A. By the different forms of *J_x_*(*y, e*), denote

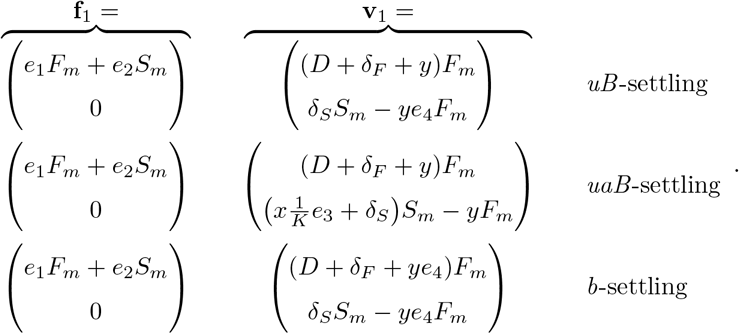

Let 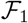 and 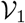 are the partial derivatives of **f**_1_ and **v**_1_ with respect to (*F_m_, S_m_*), respectively. Further, the next-generation matrix *L_x_*(*y*) is defined by 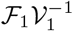. Thus, the invasion fitness proxy

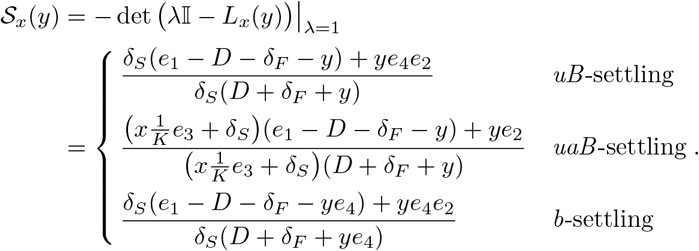

From Metz and Leimar (2011), the mutant can invade the resident if 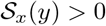, otherwise can not invade if 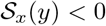. For the critical case, one can investigate the relationship of 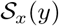, det (*J_x_*(*y, e*)) and tr(*J_x_*(*y, e*)). Further, it is easy to check the fact that – det (*J_x_*(*y, e*)) = 0 and tr(*J_x_*(*y, e*)) < 0 if 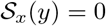. Thus, we claim that the mutant can invade the resident if 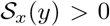, otherwise can not invade if 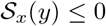.

## Appendix F The selection gradient, 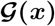

For the case of *uB*-settling, by (5), the definition of *e* = (*e*_1_, *e*_2_, *e*_3_, *e*_4_) and the resident equilibrium *E*_+_ = (*R*\*, *F*\*, *S*\*), we further have

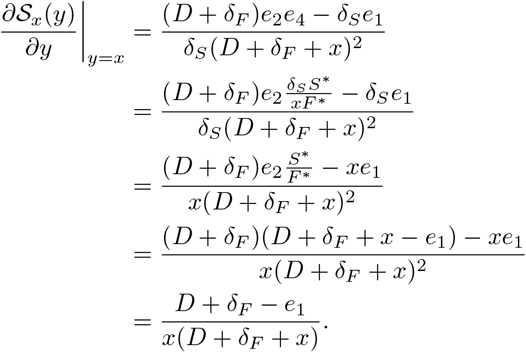

For the case of *uaB*-settling, we have

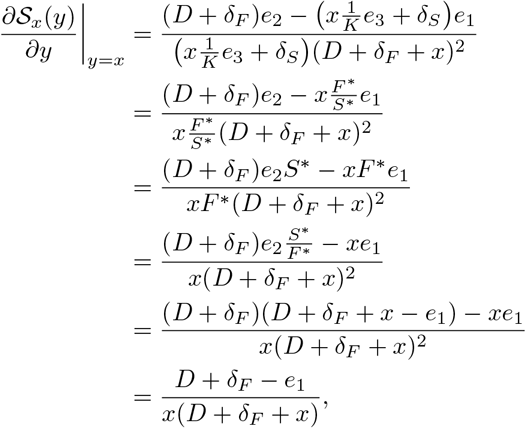

which has the same form as that of the case of *uB*-settling.

For the case of *b*-settling, we also have

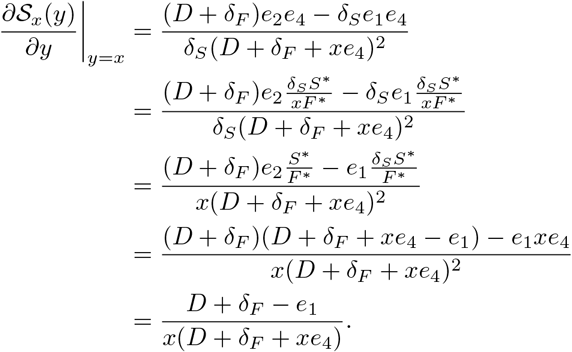

To sum up, the monomorphic selection gradient

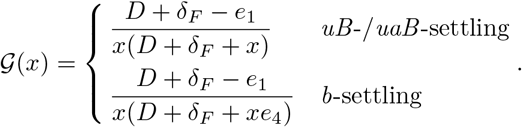

## Appendix G Existence and uniformly persistence of positive periodic solutions of the periodically forced system

The periodically forced system is the system (2) but with periodic nutrient input (7). Let 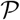 is the Poincar*é* map associated the periodically forced system, i.e., 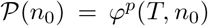 for 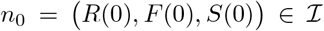, where period 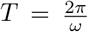 and *φ^p^*(*t, n*_0_) is the unique solution of the periodically forced system with *φ^p^*(0, *n*_0_) = *n*_0_. As shows in Appendix B, 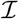 and 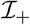 are positively invariant, 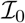 is a relative closed set in 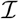. Notice that 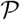 is point dissipative. Let

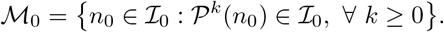

In the periodically forced system, the washout steady state 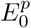 is time-varying and is *T*-periodic. Both 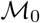 and 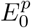 are fixed points of 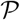 in 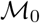.

We now claim that if 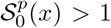, there exists *η* > 0 such that when 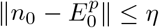 for any 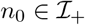, we have

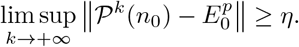

Suppose, by contradiction, that 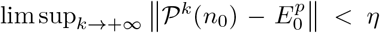 for some 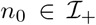. Then there exists an integer 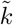 such that 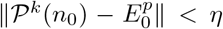 for all 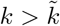. By the continuity of the solution *φ^p^*(*t, n*_0_),

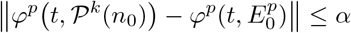

for all *t* ≥ 0 and *α* > 0. Let *t* = *kT* + *τ* where *τ* ∈ [0, *T*) and *k* is the greatest integer less than or equal to 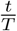. If *n*_0_ satisfies 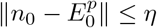, then

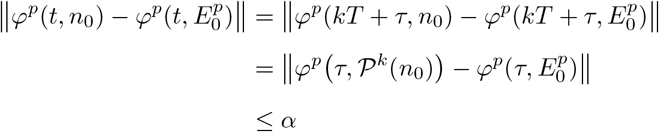

for all *t* ≥ 0. By the definition of *φ^p^*(*t*, ·) and 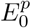, it follows that 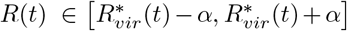 and (*F*(*t*), *S*(*t*)) ∈ [0, *α*]^2^ for all *t* ≥ 0, where 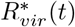 is the periodic solution of the differential equation 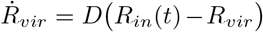. Since 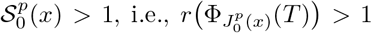, we always can choose *α* > 0 small enough such that the *α*-perturbation of 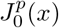, denoted by 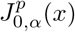, which has

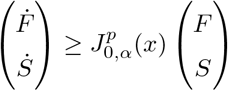

and 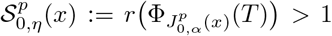 (see, e.g., Zhang and Zhao 2007). By the standard comparison principle, we have

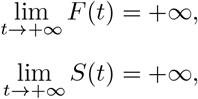

which leads to a contradiction with the dissipative of 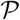. Thus, we reach the assertion.

By the above claim, it then implies that 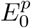 is isolated invariant in 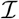, and the stable manifold of 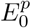 doesn’t intersect with 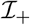. Thus, 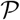 is uniformly persistent with respect to 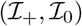 if 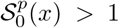. By Zhao (2003, Theorem 3.1.1), it implies that the solution of the periodically forced system with initial value 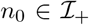 is uniformly persistent. Furthermore, by Zhao (2003, Theorem 1.3.6), 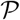 has a fixed point in 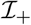. That is to say, there exists a *T*-periodic solution (*R*^*^(*t*), *F*^*^(*t*), *S*^*^(*t*)) in 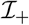. From the dynamics of the nutrients, it is clear that *R*^*^(*t*) > 0 for all *t* > 0. Thus, (*R*\*(*t*), *F*\*(*t*), *S*\*(*t*)) is a positive *T*-periodic solution.

